# Organoid Profiler: Automated, high-throughput and quantitative morphological characterization uncovers conserved longitudinal developmental kinetics in microfluidics-engineered organoids

**DOI:** 10.64898/2026.01.01.694533

**Authors:** Edgar A. Galan, Wanlong Wang, Yu Zhu, Zitian Wang, Jie Wang, Alex N. Feldman, Yongde Cai, Gan Sang, Xiaoyong Dai, Shaohua Ma

## Abstract

Three-dimensional organoid systems have emerged as transformative tools in biomedical research, yet their translation into standardized high-throughput screening platforms is impeded by stochastic fabrication variability and a lack of scalable quantitative analytics. Here, we present a unified framework integrating microfluidic droplet engineering with *Organoid Profiler*, an automated deep-phenotyping pipeline designed to capture longitudinal morphological kinetics. By processing over 10,000 longitudinally tracked images across murine lung and liver and human cerebral models, we show that this platform achieves human-level segmentation precision (*r* = 0.99) while uncovering conserved developmental dynamics obscured in manual cultures. We identify a distinct biphasic “remodeling-to-expansion” trajectory in microfluidic organoids, characterized by an initial compaction phase underpinned by the transcriptional upregulation of focal adhesion pathways, followed by exponential expansion. We further demonstrate a “shape relaxation” phenomenon where tissues converge to spherical equilibrium regardless of initial geometry, indicating intrinsic self-organization. Multi-modal profiling confirms that microfluidic organoids exhibit superior viability, homogeneity (area variability ≤ 5% upon fabrication, ≤ 15% across development) and biological fidelity, retaining physiological levels of immune and stromal niche components often lost in conventional culture. This study establishes morphological metrics as reliable, non-invasive proxies for tissue health and molecular maturity, providing a standardized foundation for quantitative organoid biology.

## Introduction

Organoid culture systems provide a physiologically relevant bridge between two-dimensional cell lines and *in vivo* models, recapitulating the multicellular architecture and functional complexity of native tissues.^1^ However, the transition of organoid technology from qualitative research tools to robust high-throughput screening platforms remains constrained by the inherent stochasticity of conventional fabrication methods.^2,3^ Manual embedding in extracellular matrix (ECM) domes frequently yields heterogeneous populations with high batch-to-batch variability in size and structure, which confounds experimental data and reduces the sensitivity of drug assays.^4^ To address this, microfluidic droplet-based technologies have been developed to generate monodisperse organoids within controlled microenvironments.^5–8^ Yet, the high-throughput generation of thousands of organoids creates a secondary bottleneck: the need for equally scalable analytical tools capable of extracting quantitative biological insights from massive image datasets.

Current analytical approaches largely rely on labor-intensive manual quantification or complex deep-learning models that require extensive annotated training datasets.^9–11^ Biological studies often focus on static endpoints, failing to capture the dynamic longitudinal kinetics—such as symmetry breaking, lumen formation, and compaction—that define healthy tissue establishment.^12,13^ Consequently, there is an unmet need for an accessible, automated framework that couples precise microfluidic fabrication with longitudinal, high-content morphometric profiling.

Here, we present *Organoid Profiler*, a robust image analysis pipeline that democratizes the quantitative characterization of organoid morphology. We integrate this software with a monodisperse microfluidic droplet generation system to establish a unified platform for high-throughput organoid engineering. By processing longitudinally tracked datasets across diverse tissue models—including primary mouse lung and liver, and human induced pluripotent stem cell (hiPSC)-derived cerebral organoids—we demonstrate that this approach achieves human-level precision while revealing conserved developmental programs distinct from manual cultures.

Specifically, we identify a conserved “remodeling-to-expansion” growth trajectory in microfluidic organoids, where initial compaction is driven by the upregulation of cell adhesion pathways. We demonstrate the platform’s capacity to tune organoid geometry, revealing a “shape relaxation” phenomenon where tissues autonomously converge to spherical equilibrium. Multi-modal validation using transcriptomics, immunofluorescence, and metabolic assays confirms that microfluidic organoids exhibit enhanced viability and biological fidelity compared to non-engineered controls, specifically retaining immune and stromal niche components. This study establishes a standardized framework for quantitative organoid research, validating morphological metrics as rigorous non-invasive proxies for molecular maturity.

## Results

### High-throughput, automated characterization of organoid morphology and growth with *Organoid Profiler*

To enable quantitative, high-throughput analysis of organoid development, we developed *Organoid Profiler*, a standardized image analysis pipeline designed to process large-scale microscopy datasets (**Fig. 1**). The workflow is compatible with brightfield and fluorescence imaging as well diverse organoid fabrication protocols, ranging from high-throughput microfluidic droplet-based cultures to traditional manual Matrigel dome embedding (**Fig. 1a–f** ).

**Fig. 1:**
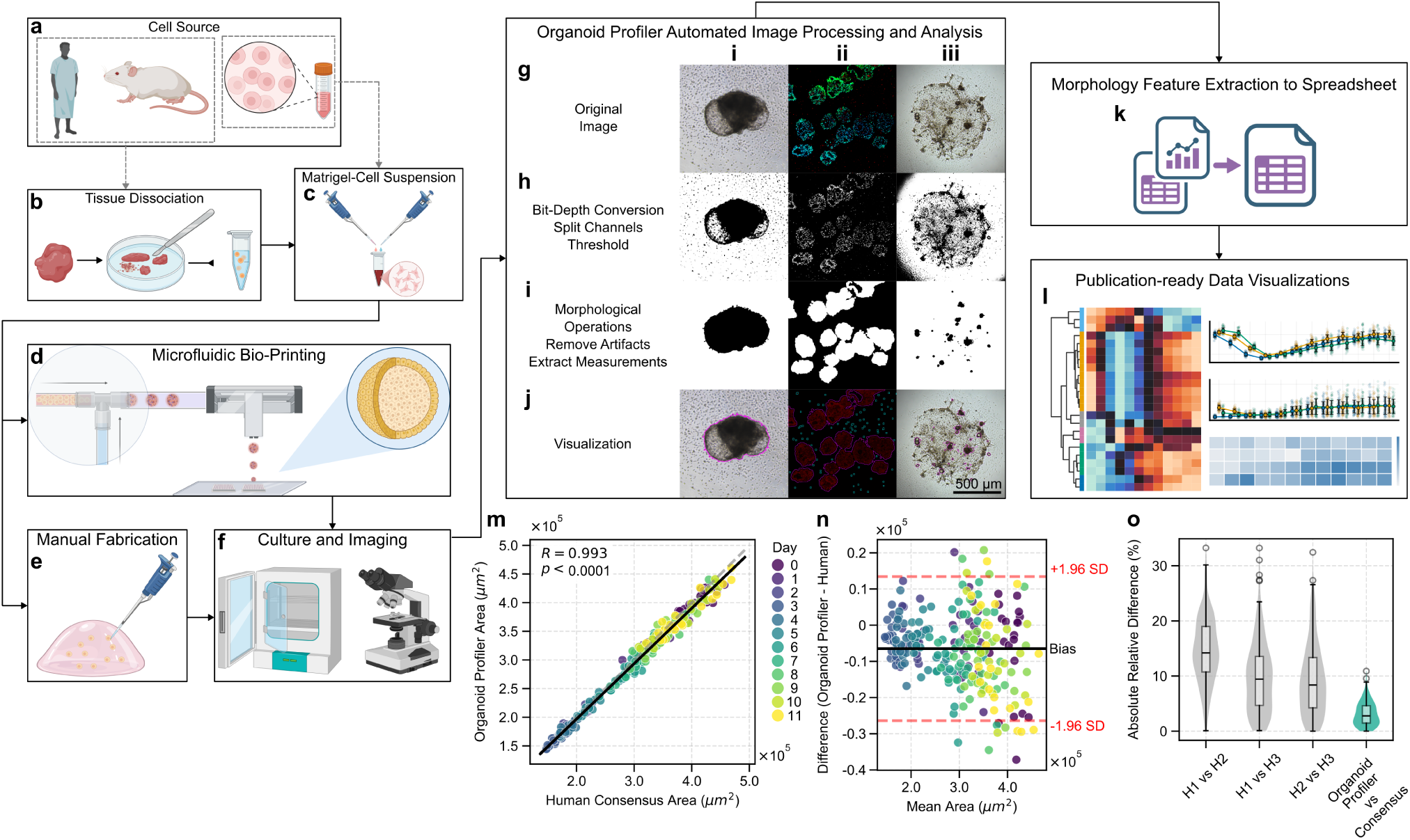
*Organoid Profiler*: High-throughput organoid morphological characterization pipeline. **a**–**f** Schematic of the organoid fabrication and imaging protocol. **a** Cell sourcing from human/mouse tissue or cell lines. **b** Tissue dissection and dissociation. **c** Suspension in Matrigel. **d** Microfluidic droplet generation for high-throughput culture. **e** Manual Matrigel dome plating. **f** Incubation and time-lapse microscopy. **g**–**j** Automated image processing workflow. Columns show representative examples of **i** a single microfluidic organoid (brightfield), **ii** immunofluorescence stained organoids, and **iii** traditional non-microfluidics organoids in a Matrigel dome (brightfield). Rows depict **g** raw input images, **h** pre-processing (bit-depth conversion, channel splitting, thresholding), **i** morphological operations (artifact removal, measurement extraction), and **j** segmentation results (magenta outlines) overlaid on original images; cyan outlines in column **ii** indicate excluded non-organoid features. Scale bar, 500 *μ*m. **k**, **l** Data output modules for **k** quantitative feature extraction (CSV) and **l** longitudinal visualization. **m**–**o** Validation of *Organoid Profiler* accuracy against manual quantification (consensus of 3 independent researchers). **m** Pearson correlation of organoid area measurements comparing *Organoid Profiler* to the manual consensus (*n* = 240 organoids, *r* = 0.99, two-sided *P* < 0.0001). **n** Bland-Altman analysis of agreement (Bias = −6, 473 *μ*m^2^; dashed lines represent 95% limits of agreement: −26, 426 to 13, 481 *μ*m^2^). **o** Violin plots comparing measurement variability between researchers (grey; H1 vs H2, H1 vs H3, H2 vs H3) and between *Organoid Profiler* and the researcher consensus (green). Source data are provided as a Source Data file.

The pipeline automates the extraction of morphological features through a series of robust image processing steps (**Fig. 1g–j**). Raw images (brightfield or fluorescence) undergo pre-processing, followed by automated thresholding to segment regions of interest. To ensure data integrity, the algorithm applies morphological filters—such as circularity and size constraints—to distinguish true organoids from debris, artifacts, or single cells, automatically excluding non-organoid features from the final dataset (**Fig. 1j**). The system outputs comprehensive morphological metrics (e.g., area, perimeter, shape factors) in a standardized format (CSV; **Fig. 1k**) and facilitates the visualization of longitudinal developmental trajectories (**Fig. 1l**).

We validated the accuracy of *Organoid Profiler* by comparing its automated area measurements against a consensus dataset derived from manual annotations by three independent researchers (*n* = 240 organoids). The automated measurements demonstrated a high degree of linearity relative to the manual consensus (Pearson’s *r* = 0.99, *P* < 0.0001; **Fig. 1m**). Bland-Altman analysis indicated high agreement with a marginal bias of −6, 473 *μ*m^2^ (2.2% of the mean organoid area) and limits of agreement spanning −26, 426 to 13, 481 *μ*m^2^ (8.9% to 4.5% of the mean organoid area, respectively) (**Fig. 1n**). Furthermore, the variability between the automated system and the manual consensus was comparable to the inter-researcher variability observed among human annotators (**Fig. 1o**), demonstrating that *Organoid Profiler* achieves human-level precision while enabling high-throughput, unbiased quantification.

To ensure reproducibility and standardize morphological nomenclature across the field, we explicitly defined the mathematical algorithms for the 25 distinct features extracted by the pipeline (**Extended Data Table 1**). These metrics are categorized into geometric descriptors (e.g., *Area*, *Perimeter*, *Feret Diameter*) to quantify size and growth; shape factors (e.g., *Circularity*, *Solidity*, *Roundness*) to assess boundary regularity and symmetry; and intensity-based statistics (e.g., *Skewness*, *Kurtosis*, *Corrected Total Intensity*) to serve as non-invasive proxies for tissue density and optical opacity. By codifying these definitions, *Organoid Profiler* eliminates the ambiguity often associated with manual qualitative assessments, allowing for the precise decoupling of size, shape, and texture dynamics. These data enable the building of explainable morphometric organoid data banks.

### Longitudinal high-content profiling reveals distinct morphological phases in microfluidics-engineered organoids

To characterize the growth dynamics and morphological evolution of microfluidics-engineered organoids, we generated monodisperse populations of primary mouse lung organoids, then we employed *Organoid Profiler* to extract and analyze multiparametric features from brightfield microscopy time-series data (**Fig. 2a–c**; **Extended Data Table 1** for feature definitions). High-throughput longitudinal tracking of three independent batches (*n* = 96 organoids per batch) revealed distinct phenotypic signatures that were categorized into six functional groups: size, contour, elongation, intensity, distribution, and other (**Fig. 2d–f** ). Unsupervised hierarchical clustering of these features identified conserved temporal patterns across all batches, highlighting the reproducibility of the microfluidic fabrication method (**Fig. 2d–f** ).

**Fig. 2:**
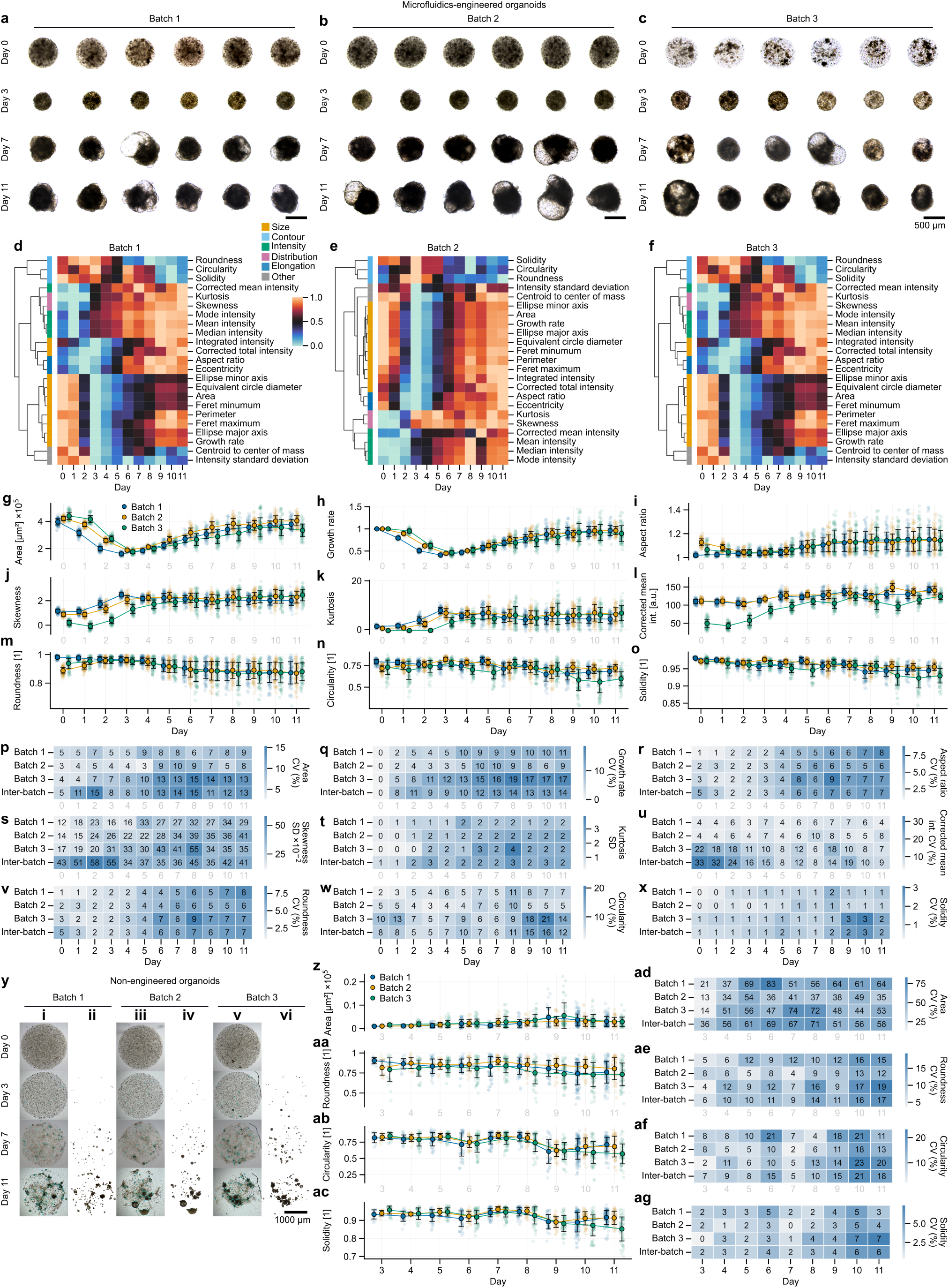
Comparative longitudinal tracking and high-content morphometric profiling of microfluidics-engineered and non-engineered lung organoids. **a–c** Representative brightfield microscopy time-series montages of three independent batches (Batch 1, Batch 2, Batch 3) of microfluidics-engineered primary mouse lung organoids. Individual organoids were tracked longitudinally, with snapshots displayed at Days 1, 3, 7, and 11 of culture. **d–f** Unsupervised hierarchical clustering (clustermaps) of morphometric features extracted from the respective organoid batches in **a–c** (*n* = 96 organoids per batch). Features were extracted using *Organoid Profiler*, normalized (z-score), and clustered to visualize morphological homogeneity and phenotypic signatures across the populations. Features are categorized into six distinct functional groups: Size (orange), physical dimensions and magnitude; Contour (light blue), smoothness and regularity of the boundary; Elongation (dark blue), deviation from a perfect circle based on elliptical fitting; Intensity (green), optical density and brightness statistics; Distribution (purple), statistical distribution of pixel intensities; and Other (gray), miscellaneous descriptors regarding internal symmetry and variance. **g–o** Longitudinal quantitative trajectories of microfluidics-engineered organoids for key morphometric features: **g** area, **h** growth rate, **i** aspect ratio, **j** skewness, **k** kurtosis, **l** corrected mean intensity, **m** roundness, **n** circularity, and **o** solidity. Data are presented as line plots of mean ± s.d. overlaid with strip plots to visualize individual organoid distribution; each dot represents one organoid. **p–x** Heatmaps quantifying variability across culture days for microfluidics-engineered batches (Batch 1, Batch 2, Batch 3) and the inter-batch combination. Coefficient of variation (CV) is displayed for **p** area, **q** growth rate, **r** aspect ratio, **u** corrected mean intensity, **v** roundness, **w** circularity, and **x** solidity. Standard deviation (s.d.) is displayed for **s** skewness and **t** kurtosis. **y** Representative images of non-engineered organoids at Days 0, 3, 7, and 11. Columns **i**, **iii**, and **v** display representative brightfield images of **i** Batch 1, **iii** Batch 2, and **v** Batch 3 organoids. Columns **ii**, **iv**, and **vi** display *Organoid Profiler* segmentation results of the images in **i**, **iii**, and **v**, respectively. Scale bar: 1000 *μ*m. **z–ac** Longitudinal quantitative trajectories of non-engineered organoids for **z** area, **aa** roundness, **ab** circularity, and **ac** solidity. Data are presented as line plots of mean ± s.d. overlaid with strip plots; each dot represents one organoid. **ad–ag** Heatmaps quantifying variability across culture days for non-engineered organoid batches (Batch 1–3) and the inter-batch combination. Coefficient of variation (CV) is displayed for **ad** area, **ae** roundness, **af** circularity, and **ag** solidity. Source data are provided as a Source Data file.

Quantitative trajectory analysis of key morphometric features (**Fig. 2g–o**) revealed a biphasic growth pattern characteristic of microfluidics-engineered organoids suspended in culture. In the initial phase (Days 0–3), organoids exhibited a “remodeling” period defined by a significant decrease in projected area (**Fig. 2g**) and growth rate (**Fig. 2h**), accompanied by an increase in roundness (**Fig. 2m**) and circularity (**Fig. 2n**). This compaction results from the reorganization of the Matrigel scaffold by the encapsulated single cells as they establish cell-cell contacts, a process distinct from the substrate-adhered growth observed in traditional dome cultures. Following this remodeling phase (Day 3 onwards), organoids entered an “expansion” phase characterized by a steady increase in size and growth rate (**Fig. 2g, h**) driven by cell proliferation.

As the organoids matured, their optical properties and shape complexity evolved. The formation of a central lumen, typically visible between Days 3 and 4 (**Fig. 2b**), coincided with distinct shifts in texture and intensity metrics. We observed a marked increase in skewness (**Fig. 2j**) and kurtosis (**Fig. 2k**) alongside a rise in corrected mean intensity (**Fig. 2l**), reflecting the transition from a transparent, low-density cell suspension to a denser, optically complex tissue structure. Furthermore, the emergence of multiple fluid-filled, alveoli-like structures protruding from the main body post-Day 3 led to a gradual deviation from spherical symmetry, quantitatively captured by increasing aspect ratio (**Fig. 2i**) and decreasing circularity and roundness (**Fig. 2m, n**). Minor fluctuations in solidity (**Fig. 2o**) were attributed to cells occasionally adhering to the organoid periphery.

To assess the tunability and homogeneity of the fabrication platform, we analyzed the inter and intra-batch variability of these features (**Fig. 2p–x**). Heatmaps of the coefficient of variation (CV) demonstrated low batch-to-batch variability for shape parameters such as roundness (**Fig. 2v**) and circularity (**Fig. 2w**), particularly during the remodeling phase, underscoring the high monodispersity achieved by the microfluidic flow-focusing technique. While size heterogeneity (Area CV) increased moderately during the expansion phase (**Fig. 2p**), the consistent trajectory of morphometric features across independent batches confirms the robustness of the platform for generating standardized organoid models.

To demonstrate the generalizability of *Organoid Profiler* to other tissue models, we analyzed the developmental trajectory of microfluidics-engineered primary mouse liver organoids (**Extended Data Fig. 1**). Longitudinal tracking of a separate batch (*n* = 192) revealed a distinct morphometric footprint compared to lung organoids. Unlike the biphasic growth pattern observed in lung cultures, liver organoids exhibited a continuous contraction phase, with a steady decrease in projected area (**Extended Data Fig. 1d**) and growth rate (**Extended Data Fig. 1e**) over 13 days. Furthermore, while lung organoids typically established a central lumen by Day 3–4, liver organoids acquired complex cystic morphologies later in culture, typically around Day 7, as evidenced by the delayed divergence in shape parameters such as solidity (**Extended Data Fig. 1l**) and circularity (**Extended Data Fig. 1k**). This capability to quantify subtle, tissue-specific developmental differences underscores the versatility of *Organoid Profiler* for diverse organoid systems.

We further extended this validation to human induced pluripotent stem cell (hiPSC)-derived cerebral organoids, which exhibit complex, multi-stage developmental kinetics distinct from primary tissue-derived models (**Extended Data Fig. 2**). Quantitative profiling evidenced a trajectory strictly governed by the differentiation protocol. During the initial Embryoid Body (EB) formation (Days 0–4), organoids displayed rapid growth, characterized by a sharp increase in projected area and growth rate (**Extended Data Fig. 2d, e**). The transition to neural induction medium on Day 5 triggered a symmetry-breaking event, quantitatively captured by a distinct spike in the variability of aspect ratio, skewness, kurtosis, and roundness by Day 8 (**Extended Data Fig. 2c, f–h**). Following transfer to Matrigel droplets (Day 9 onwards), the organoids underwent neuroepithelial budding and expansion; this developmental milestone was marked by a decrease in skewness and kurtosis but a dramatic reduction in circularity and solidity (**Extended Data Fig. 2j–l**) alongside increased heterogeneity in size and growth rate (**Extended Data Fig. 2c**). Immunofluorescence staining for the apical marker aPKC, neural progenitor marker N-cadherin, and neuronal marker TUJ1 at Day 11 confirmed the cerebral identity of the tissues (**Extended Data Fig. 2m–o**). These data demonstrate that *Organoid Profiler* is sufficiently sensitive to detect morphological inflection points induced by media transitions, facilitating the monitoring and optimization of complex differentiation protocols.

Finally, to benchmark the performance of the microfluidic platform against the current standard, we utilized *Organoid Profiler* to characterize non-engineered organoids fabricated via the manual Matrigel dome method (**Fig. 1e**). In contrast to the uniform, monodisperse populations observed in microfluidic cultures, non-engineered organoids exhibited significant stochasticity in establishment and growth across three independent batches (**Fig. 2y**). Longitudinal quantification revealed a divergent growth pattern: while microfluidics-engineered organoids displayed tight intra-batch size distributions (**Fig. 2p**), non-engineered organoids showed a progressive increase in size heterogeneity, with the coefficient of variation (CV) for projected area escalating from 36.5% at Day 3 to 58.2% by Day 11 (**Fig. 2z, ad**) compared to 5% at Day 0 and 13% by Day 11 in microfluidics-engineered organoids (**Fig. 2g, p**). Furthermore, shape descriptors indicated a failure to maintain geometric uniformity; unlike microfluidic organoids which converged towards spherical equilibrium, non-engineered organoids became increasingly irregular over time, evidenced by a decline in mean Circularity (0.84 to 0.69) and Solidity (0.94 to 0.90) (**Fig. 2aa–ac**) alongside high intra-batch variability (**Fig. 2ae–ag**). These data demonstrate that the microfluidic fabrication method effectively minimizes the “batch effect” and morphometric noise inherent to manual culture, yielding standardized organoid populations suitable for high-content screening, which *Organoid Profiler* can rigorously characterize.

### Optimization of seeding density reveals tunability and reproducibility in microfluidics-engineered organoids

To establish the optimal parameters for organoid fabrication, we utilized *Organoid Profiler* to characterize the morphological kinetics of primary mouse lung organoids generated at three distinct initial cell concentrations: low (0.2 × 10^5^cells ml^−1^), medium (1.0 × 10^5^cells ml^−1^), and high (2.5 × 10^5^cells ml^−1^). Visual inspection of longitudinal brightfield images indicated that while organoids formed in all conditions, the low-density group produced smaller and less uniform structures compared to the medium and high-density groups, which appeared morphologically indistinguishable (**Fig. 3a–c**).

**Fig. 3:**
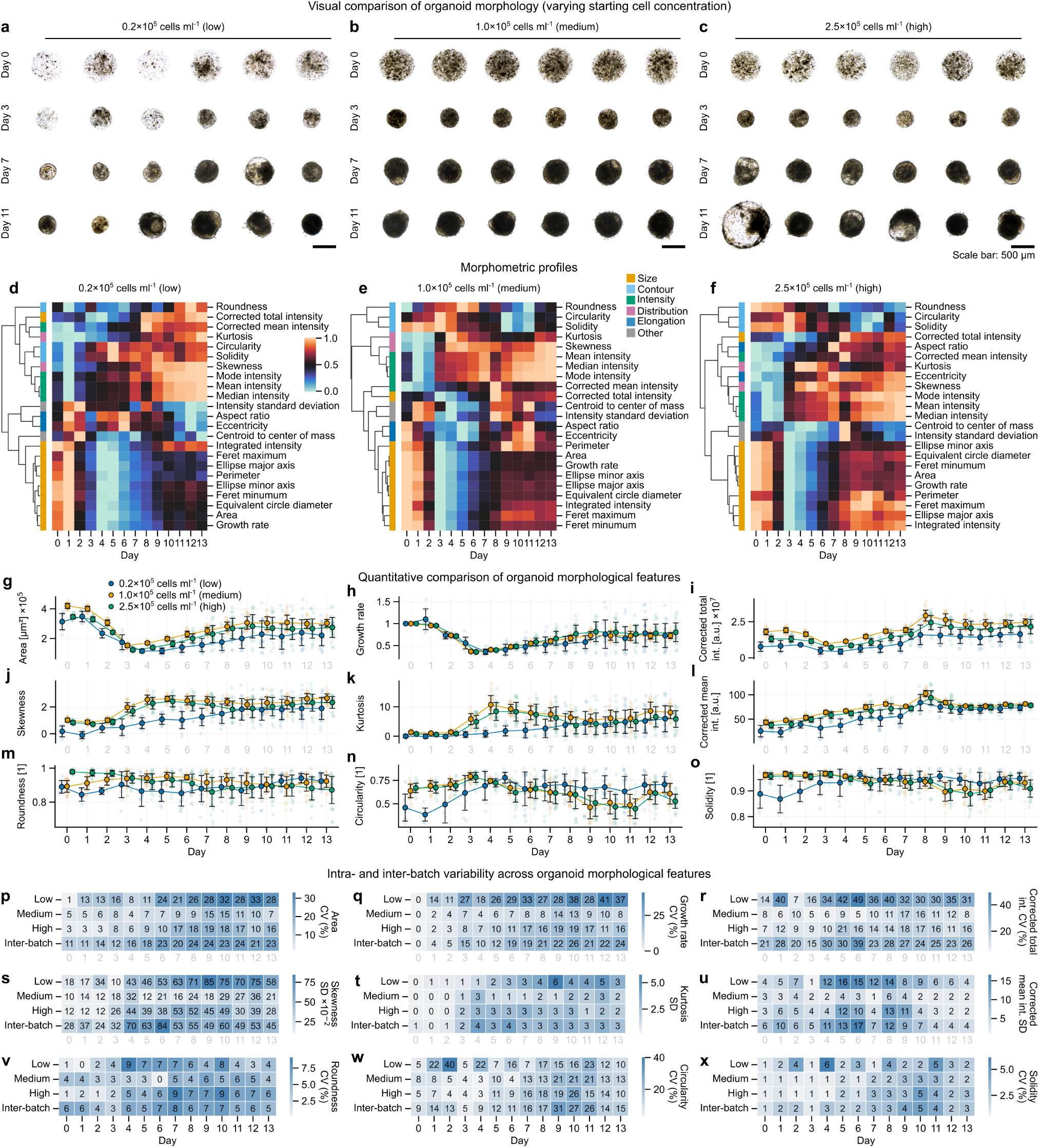
Quantitative morphometric profiling and variability analysis of microfluidics-engineered organoids across varying starting cell concentrations. **a–c** Representative brightfield images of primary mouse lung organoids generated via microfluidic droplet encapsulation and cultured in 96-well plates. Organoids were fabricated using three different initial cell concentrations: **a** low (0.2 × 10^5^ cells ml^−1^), **b** medium (1.0 × 10^5^ cells ml^−1^), and **c** high (2.5 × 10^5^ cells ml^−1^). Rows display the longitudinal development of individual organoids at Days 0, 3, 7, and 11. Images are representative of *n* = 32 organoids analyzed per condition. Scale bar: 500 *μ*m. **d–f** Longitudinal morphometric profiles of organoid development for **d** low, **e** medium, and **f** high starting cell concentrations. Heatmaps display the temporal evolution of morphological features (rows) from Day 0 to 13. Data represent the mean value of *n* = 32 organoids per condition. Features are min-max scaled (0–1) to normalize dynamic ranges and hierarchically clustered (Euclidean distance, average linkage) to reveal parameters with similar temporal kinetics. Color groups on the left indicate feature categories: Size (orange), Contour (light blue), Elongation (dark blue), Intensity (green), Distribution (purple), and Other (gray). **g–o** Quantitative comparison of organoid morphological features. Line plots represent the mean ± s.d. of *n* = 32 organoids per condition. Individual data points are shown as overlaid strip plots. Parameters plotted are: **g** Area, **h** Growth rate, **i** Corrected total intensity, **j** Skewness, **k** Kurtosis, **l** Corrected mean intensity, **m** Roundness, **n** Circularity, and **o** Solidity. Blue: Low; Orange: Medium; Green: High. **p–x** Intra and inter-batch variability across organoid morphological features. Heatmaps display the coefficient of variation (CV) or standard deviation (s.d.) for each parameter over time. Rows represent the intra-batch variability for the three starting cell concentrations (Low, Medium, High) and the inter-batch variability calculated across the combined dataset. Panels show CV for **p** Area, **q** Growth rate, **r** Corrected total intensity, **u** Corrected mean intensity, **v** Roundness, **w** Circularity, and **x** Solidity; and s.d. for **s** Skewness and **t** Kurtosis. Lighter colors indicate lower variability (higher homogeneity). Source data are provided as a Source Data file.

Hierarchical clustering of morphometric profiles confirmed that organoid development follows distinct temporal phases regardless of seeding density (**Fig. 3d–f** ). However, quantitative analysis highlighted critical differences in growth kinetics and homogeneity. While the low-concentration group exhibited stunted growth and consistently lower projected areas (Day 13 mean area ≈ 2.09 × 10^5^ *μ*m^2^; **Fig. 3g**), the medium and high-concentration groups displayed overlapping growth trajectories. Notably, increasing the cell concentration from medium to high did not yield a significant increase in final organoid size (Day 13 mean area ≈ 2.98 × 10^5^ *μ*m^2^ vs 2.78 × 10^5^ *μ*m^2^, respectively) or growth rate (**Fig. 3h**), suggesting that the medium concentration provides sufficient cellular mass for optimal organoid formation without unnecessary consumption of primary tissue.

We observed that optical properties significantly influenced morphological quantification during early culture stages. In the initial days (Day 0–3), organoids exhibited low Skewness and Kurtosis (**Fig. 3j, k**; **Extended Data Table 1**), indicative of the Gaussian-like pixel intensity distributions typical of optically transparent, loose cell aggregates. As organoids matured and densified (Day 4+), these transparency-dependent metrics increased and stabilized. Similarly, the challenge of accurately segmenting the boundaries of translucent, immature aggregates resulted in lower mean values and higher standard deviations for shape descriptors such as Roundness, Circularity, and Solidity during the first 72 hours (**Fig. 3m–o**). Following this aggregation phase, these metrics plateaued near unity, reflecting the formation of robust, quasi-spherical three-dimensional structures.

Crucially, analysis of intra- and inter-batch variability demonstrated that the medium cell concentration offers a superior balance between reproducibility and efficiency (**Fig. 3p–x**). Heatmaps of the coefficient of variation (CV) revealed that the medium concentration yielded homogeneity comparable to, and in some metrics superior to, the high-concentration group, particularly in size (5% at Day 0 and 7% by Day 13 for the medium-concentration group compared to 3% at Day 0 and 16% by Day 13 in the high-concentration group) (**Fig. 3p**) and shape regularity (**Fig. 3v–x**). In contrast, the low-concentration group showed higher stochasticity across multiple markers, reaching an Area CV of 28% by Day 13. Consequently, the 1.0 × 10^5^ cells ml^−1^ concentration was identified as the optimal condition, maximizing morphometric uniformity and growth potential while minimizing the quantity of primary cells required for fabrication.

### Microfluidic modulation of droplet geometry enables tunable organoid size and aspect ratio

Having established the optimal seeding density, we next evaluated the platform’s capacity to tune organoid morphology by modulating the physics of droplet generation. Using the optimized cell density of 1.0 × 10^5^ cells ml^−1^, we varied the flow rate ratio between the dispersed Matrigel phase and the continuous oil phase to generate droplets of three distinct volumes. This resulted in three organoid starting conditions: Small (1.9 × 10^5^ *μm*^2^), Medium (3.5 × 10^5^ *μm*^2^), and Large (1.2 × 10^6^ *μm*^2^) (**Fig. 4a–c**). Notably, the geometric constraints of the microfluidic tubing imposed a plug-like, high aspect ratio morphology on the Large droplets at generation (**Fig. 4c**), whereas the Small and Medium droplets retained a spherical geometry (**Fig. 4a, b**). This experimental design allowed us to simultaneously assess the platform’s control over size and its ability to monitor shape normalization dynamics.

**Fig. 4:**
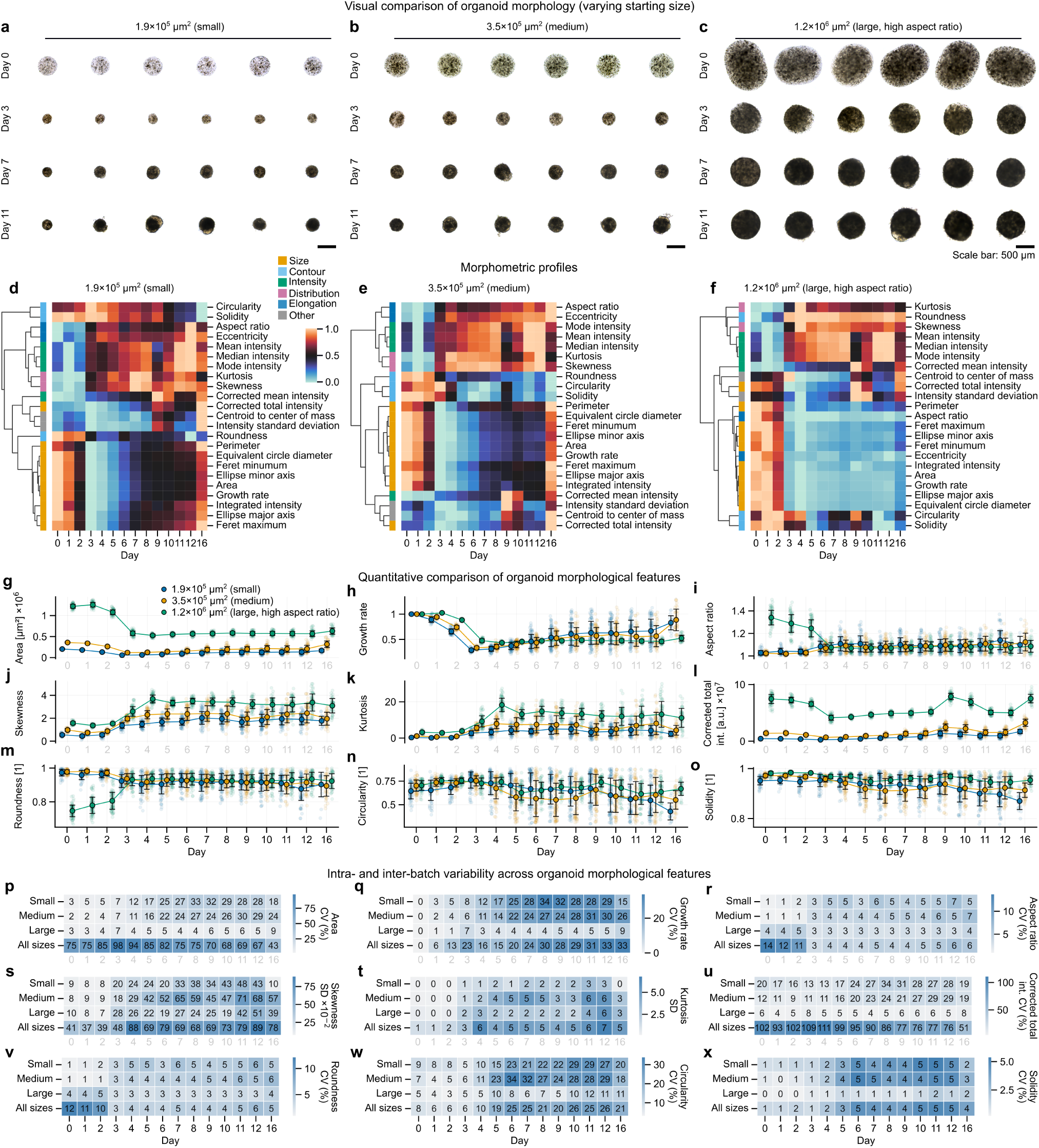
Tunability of organoid size and aspect ratio via microfluidic fabrication. **a–c** Representative brightfield images of primary mouse lung organoids derived from a single tissue sample and tracked over time (Days 0, 3, 7, and 11). Organoids were encapsulated at a constant cell density (1.0 × 10^5^ cells ml^−1^) while varying the flow rate ratio of the Matrigel phase to the oil phase to generate three distinct starting conditions: **a** Small (1.9 × 10^5^ *μm*^2^), **b** Medium (3.5×10^5^ *μm*^2^), and **c** Large (1.2×10^6^ *μm*^2^). The **c** Large group is characterized by a high initial aspect ratio compared to the circular morphology of the **a** Small and **b** Medium groups. Scale bar: 500 *μm*. **d–f** Morphometric profiles of organoids initialized at **d** Small (*n* = 109), **e** Medium (*n* = 96), and **f** Large (*n* = 83) sizes. Heatmaps display the mean values of morphological features (rows) over culture Days 0–16 (columns). Features are clustered row-wise (Euclidean distance, average linkage), while days are ordered chronologically. Data values are Min-Max normalized row-wise (scale 0–1). **g–o** Quantitative temporal evolution of selected morphological features: **g** Area, **h** Growth rate, **i** Aspect ratio, **j** Skewness, **k** Kurtosis, **l** Corrected total intensity, **m** Roundness, **n** Circularity, and **o** Solidity. Data are presented as individual organoids (dots) overlayed with line plots representing the mean ± standard deviation (s.d.). Color code: Small (blue), Medium (orange), Large (green). **p–x** Heatmaps displaying the intra-group and combined variability of organoid morphological features. Rows represent the three specific size conditions (Small, Medium, Large) and the “All sizes” condition (representing the variability of the pooled population across all three groups). Columns represent culture days. Values indicate the Coefficient of Variation (CV, %) for **p** Area, **q** Growth rate, **r** Aspect ratio, **u** Corrected total intensity, **v** Roundness, **w** Circularity, and **x** Solidity; and the standard deviation (s.d.) for **s** Skewness and **t** Kurtosis. Source data are provided as a Source Data file.

Morphometric profiling via *Organoid Profiler* revealed distinct developmental trajectories for the three conditions. Heatmap clustering indicated that while the Small and Medium groups exhibited relatively stable shape descriptors throughout culture (**Fig. 4d, e**), the Large group displayed a dynamic shift in elongation parameters during the first week (**Fig. 4f** ). Quantitative trajectory analysis confirmed that the high initial aspect ratio (∼ 1.4) of the Large group decreased rapidly, converging with the spherical morphology (aspect ratio ≈ 1.0) of the Small and Medium groups by Day 5 (**Fig. 4i**). This morphological relaxation was corroborated by synchronous increases in circularity (**Fig. 4n**) and roundness (**Fig. 4m**), demonstrating the intrinsic capacity of the organoids to self-organize into minimum-energy spherical shapes despite initial geometric deformation.

While shape parameters converged, size differences were maintained throughout the 16-day culture period. The Area trajectories (**Fig. 4g**) remained distinct and parallel, indicating that initial droplet size effectively dictates final organoid volume without compromising viability or growth potential. Importantly, the standard deviation (shaded regions) for all morphological features remained tight within each condition, suggesting high intra-group homogeneity.

To quantify this homogeneity, we calculated the coefficient of variation (CV) for each feature across the three distinct size groups. The microfluidic-controlled groups maintained low variability, though specific size-dependent trends were observed. While the Small and Medium conditions exhibited Area CVs ranging from approximately 2% to 33% over the 16-day culture period, the Large group displayed superior homogeneity, with Area CVs consistently remaining below 10% (ranging from 3.3% to 9.2%). Interestingly, shape descriptors such as Roundness and Solidity showed consistently low variability across all conditions regardless of size. Roundness CVs generally remained below 7% across all groups, while Solidity CVs were maintained at or below approximately 3% throughout the culture duration. This data confirms that while size is a tunable parameter defined by flow rates, spherical shape is an intrinsic equilibrium state of the biological system. Collectively, these results demonstrate that the platform can precisely tune organoid size while ensuring high batch homogeneity, decoupling physical dimensions from biological viability.

### Distinct morphological remodeling is accompanied by structural maturation in microfluidics-engineered organoids

To demonstrate the capability of *Organoid Profiler* to extract quantitative biological insights from large datasets in an unbiased, automated manner, we performed a longitudinal characterization of primary mouse lung organoids cultured on our microfluidic platform compared to non-engineered controls (**Fig. 5**).

**Fig. 5:**
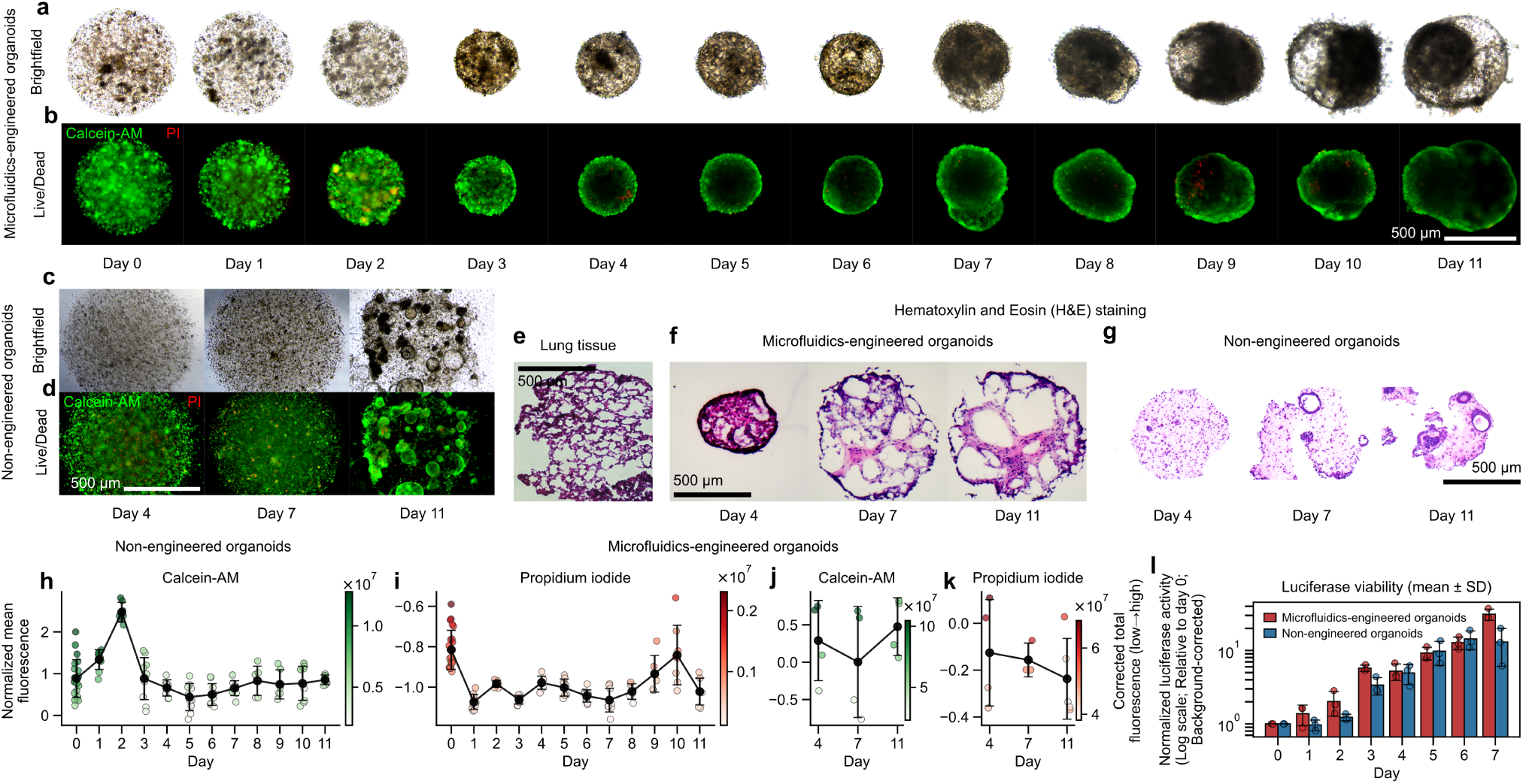
Viability, growth and morphology of microfluidics-engineered versus non-engineered organoids. **a**, **c** Representative brightfield images of **a** microfluidics-engineered organoids cultured from Day 0 to Day 11 and **c** non-engineered organoids at Days 4, 7, and 11. **b**, **d** Live/Dead fluorescence images of the organoids in **a** and **c**, respectively. Organoids were stained with Calcein-AM (live, green) and Propidium Iodide (PI) (dead, red). Scale bars, 500 *μ*m. **e**–**g** Hematoxylin and Eosin (H&E) staining sections of **e** native mouse lung tissue, **f** microfluidics-engineered organoids, and **g** non-engineered organoids at Days 4, 7, and 11, showing the preservation of hollow, cyst-like alveolar structures in microfluidic cultures compared to disorganized aggregates in non-engineered controls. Scale bars, 500 *μ*m. **h**–**k** Quantitative analysis of fluorescence intensity for Calcein-AM (live) and PI (dead) signals. **h**, **i** Microfluidics-engineered organoids (*n* = 6–24 independent organoids per time point; each dot represents one organoid). **j**, **k** Non-engineered organoids (*n* = 4–5 independent samples per time point; each sample represents an image field containing multiple organoids in a Matrigel dome). Data are presented as mean ± s.d. of the normalized mean fluorescence (left y-axis). Normalization was performed using a global Z-score calculated from the pooled dataset of both conditions to enable direct comparison of signal magnitudes. Individual data points colored according to their corrected total fluorescence intensity (color bars). **l** Longitudinal viability assessment measuring normalized luciferase activity (relative to Day 0, background-corrected). Data are presented as mean ± s.d. (*n* = 3 independent biological replicates). Source data are provided as a Source Data file.

Qualitative assessment of brightfield and live/dead fluorescence images indicated that both culture methods supported high cell viability, with minimal Propidium Iodide (PI) retention observed in either group throughout the 11-day culture period (**Fig. 5a–d**). However, the two groups exhibited distinct morphological trajectories. Microfluidics-engineered organoids developed into large, unified cystic structures characterized by multiple internal alveolar compartments encapsulated within a continuous outer lumen (**Fig. 5a**). This architecture, which preserves cell-cell contact within a shared luminal microenvironment, potentially enhances paracrine signaling and recapitulates the multi-alveolar organization of lung tissue more faithfully than non-engineered controls. In contrast, non-engineered organoids typically grew as smaller, independent cystic structures surrounded by scattered individual cells (**Fig. 5c**). Histological analysis (H&E) confirmed these observations, showing that microfluidics-engineered organoids maintained a structured, multi-compartmentalized geometry (**Fig. 5f** ) resembling native lung architecture (**Fig. 5e**), whereas non-engineered controls appeared as disjointed aggregates (**Fig. 5g**).

We employed *Organoid Profiler* to quantify these fluorescence dynamics in a standardized, high-throughput manner. To enable a direct quantitative comparison between the two culture methods, we applied a global Z-score normalization across the pooled dataset of both conditions (see **Methods**). This analysis revealed that microfluidics-engineered organoids maintained significantly higher viability than their non-engineered counterparts (**Fig. 5h–k**). While non-engineered organoids displayed lower and highly variable fluorescence intensity, consistent with the heterogeneity of the culture (**Fig. 5j, k**), microfluidics-engineered organoids exhibited a robust fluorescence trajectory (**Fig. 5h, i**).

In microfluidics-engineered organoids, *Organoid Profiler* revealed a specific intensity trajectory linked to structural remodeling (**Fig. 5h, i**). Mean Calcein-AM fluorescence increased between Day 0 and Day 2, reflecting initial proliferation and remodeling. Subsequently, mean fluorescence intensity stabilized or decreased slightly, a trend we attribute to the physical properties of the mature organoids rather than a loss of viability. As microfluidics-engineered organoids compacted and formed a sealed central lumen (Days 3–4), dye penetration became restricted to the organoid periphery, leaving the core dark due to the large volume and density of the structure. This remodeling phase—transitioning from a loose aggregate to a compact, lumenized cyst—was further evidenced by the spatial distribution of fluorescence in **Fig. 5b**, where signal is distributed throughout the volume at Day 0 but becomes circumferentially localized by Day 4. Importantly, PI levels remained low and stable after the initial culture establishment (**Fig. 5i**; Z-scores ≈ −1.0), confirming the effective clearance of initial debris and the establishment of a healthy tissue, and that the reduction in mean live signal was a geometric artifact of lumen formation and size expansion, not cell death. In contrast, non-engineered organoids showed higher relative PI levels (**Fig. 5k**; Z-scores ≈ −0.2), suggesting a lower overall health profile.

These fluorescence-based viability trends were corroborated by longitudinal metabolic analysis. Luciferase activity in microfluidics-engineered organoids peaked around Day 3—coinciding with the compaction phase—before establishing a robust exponential growth trajectory over the subsequent days (**Fig. 5l**, log scale). This metabolic profile indicates that the microfluidic platform supports highly reproducible growth (*n* = 3 independent biological replicates) and structural maturation.

Collectively, these data demonstrate that microfluidics-engineered organoids form unified, viable, and structurally complex tissues, and that *Organoid Profiler* effectively captures the quantitative signatures of this morphological maturation.

### Transcriptomic and immunofluorescence profiling reveals enhanced lineage fidelity and immune retention in microfluidics-engineered organoids

To rigorously benchmark the cellular composition and developmental trajectory of our microfluidics-engineered organoids against native tissue and conventional non-engineered organoids, we performed multi-modal characterization using bulk RNA-sequencing (RNA-seq), quantitative flow cytometry, and immunofluorescence (IF) imaging (**Fig. 6**).

**Fig. 6:**
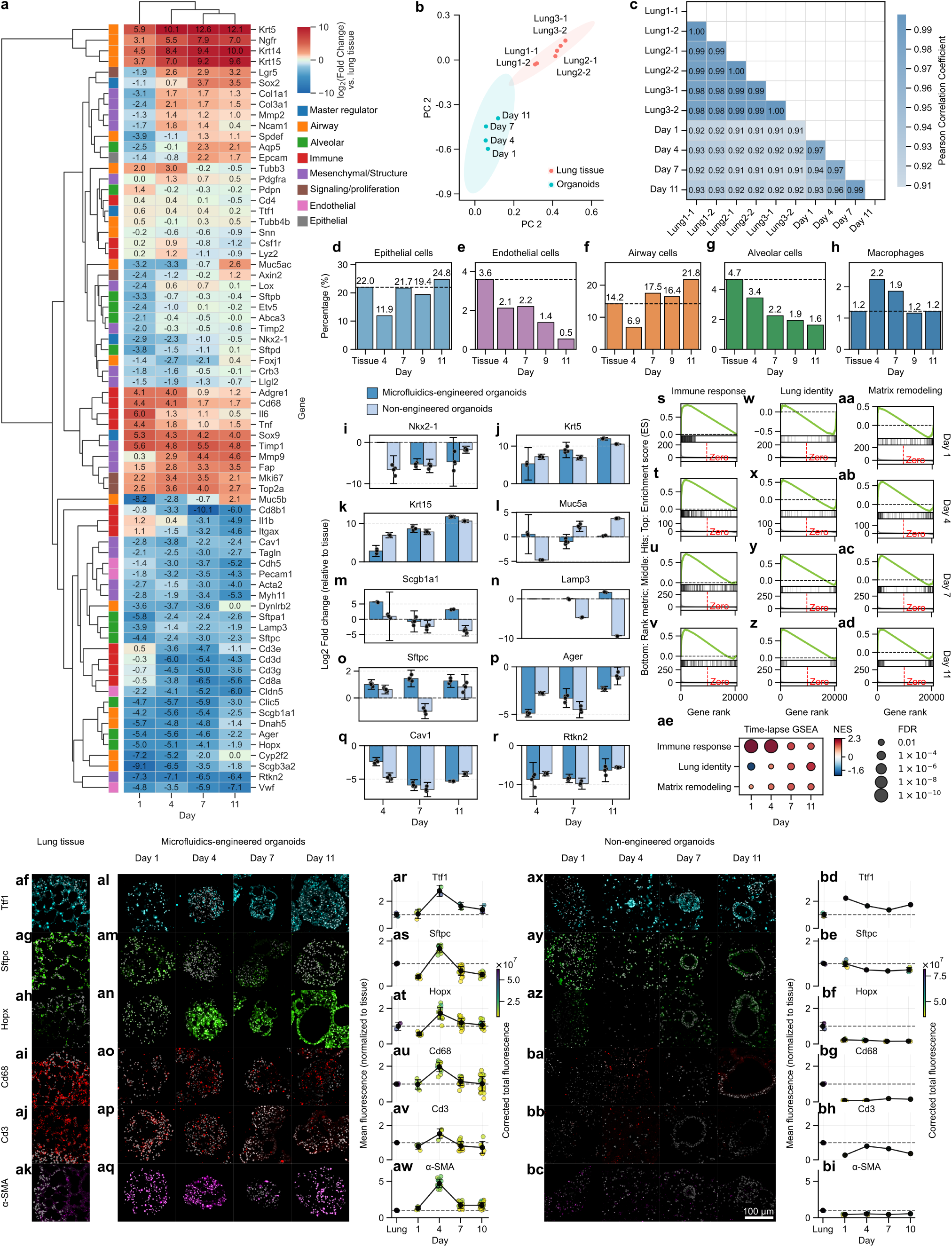
Transcriptomic and immunofluorescence characterization of organoid differentiation kinetics and homogeneity. **a** Clustered heatmap showing the differential expression (log2 fold change) of key cell type marker genes in primary mouse lung organoids at Day 1, 4, 7, and 11 relative to native mouse lung tissue controls. Genes are grouped by lineage and functional identity. The color scale indicates upregulation (red) or downregulation (blue). **b** Principal Component Analysis (PCA) of global gene expression profiles comparing primary mouse lung organoids (Day 1, 4, 7, 11) and native mouse lung tissue. PC1 and PC2 explain 37.3% and 17.1% of the variance, respectively, illustrating the separation between organoid developmental stages and native tissue. **c** Pearson correlation matrix of global gene expression (TPM) comparing biological replicates of primary mouse lung organoids and native mouse lung tissue. The color intensity represents the Pearson correlation coefficient (*r*), indicating the degree of transcriptomic similarity. **d–h** Quantitative flow cytometry analysis of cell composition in microfluidics-engineered organoids during differentiation (Days 4, 7, 9, and 11) versus native lung tissue. Bar plots display the percentage of **d** Epithelial cells, **e** Endothelial cells, **f** Airway cells, **g** Alveolar cells, and **h** Macrophages within the total viable cell population. The dashed horizontal line represents the baseline proportion measured in the native lung tissue post-digestion (Day 0) prior to organoid fabrication. Cell populations were identified using the following markers: Epithelial (CD45^−^CD31^−^CD326^+^), Endothelial (CD45^−^CD31^+^CD326^−^), Alveolar (CD45^−^CD326^+^MHCII^+^), Airway (CD45^−^CD326^+^MHCII^−^), and Macrophages (CD45^+^CD11b^+^F4/80^+^). Data represent measurements from pooled biological replicates (*n* = 3 mice) per time point. **i–r** Quantitative PCR (qPCR) analysis of lung lineage marker expression in microfluidics-engineered organoids versus non-engineered organoids at Days 4, 7, and 11 of differentiation. Panels show Log2 fold change relative to native mouse lung tissue (Day 0, dashed baseline) for **i** *Nkx2-1*, **j** *Krt5*, **k** *Krt15*, **l** *Muc5a*, **m** *Scgb1a1*, **n** *Lamp3*, **o** *Sftpc*, **p** *Ager*, **q** *Cav1*, and **r** *Rtkn2*. Data are presented as mean ±95% confidence interval (CI) with individual biological replicates (*n* = 3). Statistical analysis was performed using one-way ANOVA followed by Tukey’s multiple comparison test. **s–ad** Gene Set Enrichment Analysis (GSEA) plots showing the enrichment of **s–v** Immune response, **w–z** Lung identity, and **aa–ad** Matrix remodeling gene signatures in microfluidics-engineered primary mouse lung organoids relative to native mouse lung tissue at Days 1, 4, 7, and 11. Curves above the zero line indicate positive enrichment in the organoid phenotype. **ae** Summary dot plot of GSEA results across all time points and gene sets. The dot size represents the percentage of genes in the leading edge, and color intensity represents the Normalized Enrichment Score (NES). All displayed enrichments are statistically significant (FDR *q* < 0.05). **af–bi** Immunofluorescence analysis of lineage markers in organoids and native lung tissue. Representative images of **af–ak** native mouse lung tissue (reference), **al–aq** microfluidics-engineered organoids, and **ax–bc** non-engineered organoids stained for Ttf1 (cyan), Sftpc (green), Hopx (green), Cd68 (red), Cd3 (red), and α-SMA (magenta). Nuclei are counterstained with DAPI (gray). **ar–aw** Quantification of fluorescence intensity for **ar** Ttf1, **as** Sftpc, **at** Hopx, **au** Cd68, **av** Cd3, and **aw** α-SMA in microfluidics-engineered organoids relative to native lung tissue. **bd–bi** Quantification of fluorescence intensity for **bd** Ttf1, **be** Sftpc, **bf** Hopx, **bg** Cd68, **bh** Cd3, and **bi** α-SMA in non-engineered organoids relative to native lung tissue. Data are presented as fold change of corrected mean fluorescence intensity normalized to tissue baseline (dashed line). Source data are provided as a Source Data file.

Global transcriptomic analysis of microfluidics-engineered organoids over a differentiation time course (Days 1, 4, 7, and 11) revealed a progressive maturation trajectory. Principal Component Analysis (PCA) demonstrated a clear developmental shift along PC1 (37.3% variance), with Day 11 organoids moving toward the native lung tissue profile (**Fig. 6b**). This concordance was further corroborated by high Pearson correlation coefficients (*r* > 0.9) between organoid replicates and native tissue (**Fig. 6c**), indicating high reproducibility and transcriptomic similarity. Differential expression analysis highlighted the dynamic regulation of lineage-specific markers (**Fig. 6a**). While some alveolar markers (e.g., *Sftpc*, *Hopx*) showed lower transcript abundance relative to whole lung tissue (Log2FC ≈ −2.3 and −1.9, respectively; *P* < 0.001), we observed strong expression of basal cell markers (*Krt5*, Log2FC ≈ 12.1) and proliferation markers (*Mki67*, Log2FC ≈ 2.1), consistent with an active regenerative state.

We next assessed the cellular heterogeneity of microfluidics-engineered organoids using flow cytometry (**Fig. 6d–h** and **Extended Data Fig. 3**). Microfluidics-engineered organoids successfully retained major lung cell populations throughout culture. By Day 11, the proportion of epithelial cells (24.8%) and airway cells (21.8%) exceeded baseline tissue digests (22.0% and 14.2%, respectively), while macrophages were remarkably maintained at physiologic levels (∼ 1.2%) (**Fig. 6h**), a feature often lost in conventional organoid systems. Gene Set Enrichment Analysis (GSEA) further confirmed the functional maturation of microfluidics-engineered organoids, showing significant enrichment (FDR < 0.01) of “Immune response” (**Supplementary Data 5**), “Matrix remodeling” (**Supplementary Data 6**) and “Lung identity” (**Supplementary Data 7**) signatures at Day 11 compared to Day 1 (**Fig. 6s–ae**), validating the presence of a complex, multicellular niche.

Finally, we directly compared the protein-level expression of key lineage markers between microfluidics-engineered organoids and non-engineered organoids using quantitative immunofluorescence (**Fig. 6af–bi**). While both methods generated organoids expressing core lung transcription factors (*Nkx2-1*), microfluidics-engineered organoids demonstrated superior retention of structural and immune components. Quantification of fluorescence intensity normalized to native tissue (see **Methods**) revealed that microfluidics-engineered organoids maintained physiologic levels of the AT1 marker Hopx (1.09-fold vs. 0.16-fold in non-engineered organoids) and the macrophage marker Cd68 (1.01-fold vs. 0.15-fold in non-engineered organoids). Furthermore, microfluidics-engineered organoids exhibited a 3-fold higher retention of α-SMA^+^ mesenchymal cells (1.71-fold vs. 0.55-fold) and a 2-fold higher retention of Cd3^+^ T cells (0.72-fold vs. 0.36-fold) compared to non-engineered organoids. This enhanced preservation of the native stromal and immune niche in microfluidics-engineered organoids underscores the advantages of the microfluidic assembly approach in recapitulating the complex cellular microenvironment of the lung.

### Transcriptomic profiling confirms that organoid morphological maturation is underpinned by specific proliferation, adhesion and differentiation programs

To comprehensively characterize the molecular phenotype of the microfluidic-engineered organoids and validate that the morphological dynamics captured by *Organoid Profiler*—specifically the transition from loose aggregation to compaction and subsequent lumenized growth— we analyzed bulk RNA sequencing at four distinct timepoints (Days 1, 4, 7, and 11) and compared the organoids’ transcriptomic profiles to native mouse lung tissue (**Fig. 7**).

**Fig. 7:**
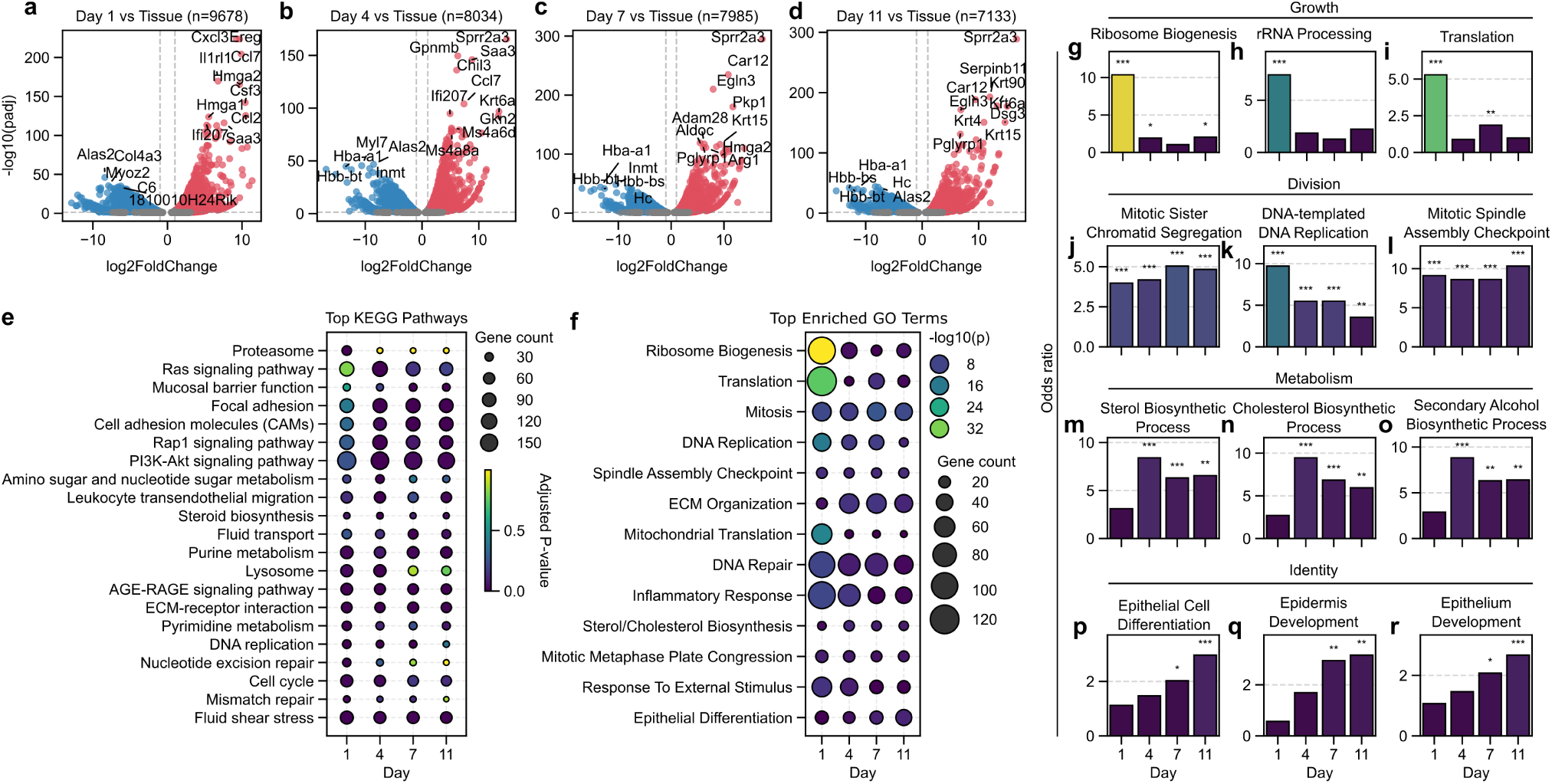
Transcriptomic characterization of organoid differentiation versus native tissue. **a–d** Volcano plots illustrating differential gene expression analysis between microfluidic primary mouse lung organoids and native mouse lung tissue at **a** Day 1, **b** Day 4, **c** Day 7, and **d** Day 11 of culture. Colored points represent significantly differentially expressed genes (log_2_ fold change > 1 or < −1; adjusted *P* < 0.05). Red points indicate genes upregulated in organoids, while blue points indicate genes downregulated in organoids relative to native tissue. Grey points indicate non-significant genes. Top differentially expressed genes are labeled in each panel. **e** Dot plot showing the enrichment of selected top KEGG pathways in organoids compared to native tissue across timepoints (Days 1, 4, 7, and 11). The size of the dots corresponds to the number of differentially expressed genes enriched in each pathway, and the color gradient indicates the significance of enrichment (adjusted *P* value). **f** Dot plot showing the enrichment of selected Gene Ontology (GO) biological process terms for genes upregulated in organoids versus native tissue across timepoints. Terms were filtered to remove redundancy; dot size represents the number of upregulated genes associated with each term, and color indicates statistical significance (adjusted *P* value). **g**–**r** Bar plots displaying the enrichment (odds ratio) of representative GO biological process terms associated with genes upregulated in organoids across culture timepoints (Days 1, 4, 7, and 11). Source data are provided as a Source Data file.

Differential gene expression analysis revealed a dynamic transcriptional landscape during organoid development that mirrors the morphological phases observed in **Fig. 2** and **Fig. 5**. At Day 1, corresponding to the initial aggregation phase where organoids appear as loose cellular clusters, we observed the upregulation of genes associated with early growth response and inflammation, such as *Ereg*, *Cxcl3*, and *Ccl7* (**Fig. 7a**). This early upregulation likely represents a transient adaptation to the in vitro microenvironment and an initial wound-healing response often seen in primary culture establishment.

As the organoids entered the morphological compaction and lumenization phase (Days 4–11), the gene expression profile shifted towards structural maturation and epithelial differentiation (**Fig. 7b–d**). We observed the strong and sustained upregulation of genes essential for epithelial integrity and barrier function, including *Sprr2a3* (Small Proline Rich Protein 2A3), *Gpnmb*, and *Saa3*, alongside cytoskeletal markers such as *Krt6a* and *Krt90*. Notably, *Car12* (Carbonic Anhydrase 12) was among the most highly upregulated genes from Day 7 onwards (**Fig. 7c, d**). Given the role of carbonic anhydrases in ion transport and pH regulation, this molecular feature is consistent with the physiological requirements of maintaining the expanding fluid-filled lumens visualized in **Fig. 5**.

Functional enrichment analysis (KEGG and GO) further bridged the morphological and molecular phenotypes. The early “metabolic startup” required to support the subsequent exponential metabolic growth described in **Fig. 5l** was evidenced by the enrichment of terms related to protein synthesis (“Ribosome”, “Translation”) at Day 1 (**Fig. 7e, f** ). We noted that while standard KEGG analysis identified broad disease-related terms often associated with high proliferation rates or immune activity (**Extended Data Fig. 4e–h**), filtering for biological relevance highlighted the specific metabolic and structural programs driving organoid formation (**Fig. 7e** and **Extended Data Fig. 4a–d**). Subsequently, the morphological stabilization and acquisition of the spherical shapes observed in **Fig. 2** were molecularly supported by the progressive enrichment of “ECM-receptor interaction”, “Focal adhesion”, and “Cell adhesion molecules (CAMs)” pathways from Day 4 to Day 11 (**Fig. 7e** and **Extended Data Fig. 4b–d**). These findings align with previous mechanobiological studies indicating that morphological perturbations directly enrich genes associated with integrins and Wnt signaling pathways.^14^ Consistent with this active mechanotransductive profile, we even observed the emergence of spontaneous, rhythmic contractions within the organoid structures by Day 7 (**Extended Data Video 1**). Furthermore, the interplay between morphology and transcriptional identity is consistent with findings in pancreatic ductal adenocarcinoma models, where TGF*β* signaling was identified as a master regulator of branching morphogenesis and EMT plasticity.^15^ Our data suggests that the “remodeling” phase detected by *Organoid Profiler* represents a critical window where these mechanotransductive and adhesive programs are established to define subsequent tissue architecture.

Crucially, while *Organoid Profiler* quantified a rapid increase in projected area (growth) during the later culture phases (**Fig. 2**), the transcriptomic data confirmed that this was driven by active proliferation. Gene Ontology (GO) biological process analysis (**Fig. 7f** ) showed that while standard results were dominated by generic biosynthetic and housekeeping processes reflecting high cellular activity (**Extended Data Fig. 5e–h**), the filtered analysis resolved the specific developmental trajectory (**Extended Data Fig. 5a–d**). Day 1 organoids were enriched for terms related to protein synthesis (“Ribosome biogenesis”, “Translation”), while terms associated with tissue organization (“Extracellular matrix organization”, “Epithelial differentiation”) progressively increased in magnitude (**Fig. 7f** and **Extended Data Fig. 5b–d**). Detailed tracking of individual GO terms (**Fig. 7g–r**) demonstrated that while proliferative signals remained robust throughout culture with upregulation of cell cycle regulators such as “Mitotic sister chromatid segregation”, “DNA-templated DNA replication” and “Mitotic spindle assembly checkpoint” (**Fig. 7j–l**), terms related to metabolism (“Sterol biosynthetic process”, “Cholesterol biosynthetic process” and “Secondary alcohol biosynthetic process”) dominated by Day 4 (**Fig. 7m–o**), and lung identity-related terms (“Epithelial cell differentiation”, “Epidermis development” and “Epithelium development”) were most prominent by Day 11 (**Fig. 7p–r**). Collectively, these data indicate that microfluidic-engineered organoids dynamic morphological and transcriptomic trajectories are interconnected through robust proliferation and complex structural remodeling programs that drive the acquisition of their tissue-like epithelial architecture.

## Discussion

The translation of organoid technology from a bespoke research tool to a standardized platform for high-throughput drug screening and regenerative medicine has been hindered by two persistent challenges: the intrinsic heterogeneity of current fabrication methods and the lack of scalable, quantitative analytical frameworks.^16,17^ In this study, we addressed these bottlenecks by integrating microfluidic droplet generation with *Organoid Profiler*, an automated image analysis pipeline. This combinatorial approach not only achieved high-throughput generation of monodisperse organoids but also uncovered conserved longitudinal developmental kinetics—specifically a biphasic “remodeling-to-expansion” trajectory—that serves as a quantitative signature of healthy organoid establishment.

While previous image analysis tools have often required complex model training or focused on specific organoid models,^9–11^ *Organoid Profiler* was designed to democratize high-content analysis by capturing the holistic morphological evolution of tissues in culture. By leveraging robust morphological operations rather than heavy computational models, the pipeline extracts 25 metrics without requiring specialized hardware (e.g., GPUs) or algorithm training. This “zero-barrier” approach—accessible via a web interface—enabled us to process a massive dataset of over 10,000 longitudinally tracked images across 12 independent batches and diverse tissue types (lung, liver, cerebral), characterizing distinct developmental programs, from the continuous contraction of liver cysts to the symmetry-breaking events of neuroectodermal differentiation. Validating the system against manual consensus (*r* = 0.99) demonstrated that this automated profiling achieves human-level precision while enabling a scale of analysis that would be prohibitive with manual annotation.

A key finding of this work is the characterization of the “morphometric noise” inherent to manual culture and its suppression via microfluidics. Non-engineered organoids exhibited stochastic growth trajectories and increasing heterogeneity over time (Area CV > 50%, compared to Area CV ≤ 15% in microfluidics-engineered organoids across all timepoints). Beyond biological variance, we identified critical physical factors contributing to this noise: non-engineered organoids embedded in Matrigel domes frequently overlap in the Z-axis and adhere to the substrate, complicating segmentation. In contrast, microfluidics-engineered organoids remain suspended in a single focal plane with minimal background debris.

This physical constraint not only facilitates precise imaging and manipulation but also ensures temporal synchronization of the culture. As demonstrated by Chiaradia et al., when tissue architecture is perturbed—for instance, by dissociation or encapsulation that disrupts cell positioning—it results in aberrant temporal progression where cells become intermingled in both space and time.^18^ By maintaining high monodispersity and structural uniformity, our microfluidic platform likely minimizes this “scrambled” fate progression, ensuring that the morphological metrics extracted by *Organoid Profiler* reflect synchronized developmental states rather than artifactual noise.

Microfluidics-engineered organoids also maintained high monodispersity and structural uniformity. This homogeneity allowed us to resolve subtle kinetic phases that are otherwise obscured by population variance in manual cultures. We identified a distinct “remodeling phase” (Days 0–3) in microfluidics-engineered organoids characterized by compaction and roundness stabilization. Transcriptomic analysis revealed that this morphological compaction is underpinned by specific molecular programs, transitioning from an initial metabolic startup to the upregulation of cell adhesion and focal adhesion pathways. This structure-function linkage suggests that the standardized metrics extracted by *Organoid Profiler* can serve as non-invasive proxies for complex molecular states.

Furthermore, we demonstrated that the physical parameters of the niche—seeding density and droplet geometry—are tunable. We optimized seeding density to conserve primary tissue and observed a “shape relaxation” phenomenon where elongated organoids converged to spherical equilibrium, underscoring the dominance of biological self-organization over initial geometry. The ability of *Organoid Profiler* to track this “shape relaxation” is biologically significant. While our platform demonstrates the intrinsic tendency of lung organoids to self-organize into minimum-energy spherical shapes, other studies utilizing non-deformable agarose microwells have shown that maintaining non-spherical geometries can fundamentally alter developmental trajectories. For example, Sen et al. demonstrated that imposing “butterfly” or “peanut” shapes accelerated neurodevelopmental kinetics and upregulated gene expression related to mechanotransduction pathways, such as Integrin and Wnt signaling, compared to spherical controls.^14^ By providing high-frequency longitudinal tracking of aspect ratio and circularity, *Organoid Profiler* enables researchers to distinguish between natural self-organization (relaxation) and geometry-induced lineage biases, a critical capability for precise tissue engineering.

Our multi-modal characterization further revealed that microfluidics-engineered organoids possess superior biological fidelity. Microfluidics-engineered organoids developed into unified, lumenized structures with complex alveolar compartmentalization, contrasting with the disorganized aggregates of manual cultures. Significantly, flow cytometry and immunofluorescence confirmed the physiological retention of macrophages, fibroblasts, and endothelial cells in microfluidics-engineered organoids. The loss of immune components is a well-documented limitation of standard organoid protocols; their preservation likely results from the rapid encapsulation of the total tissue digest within a confined, autocrine-rich microenvironment, positioning the microfluidics-engineered organoid platform as a superior model for investigating immune-epithelial interactions and inflammatory responses.

We acknowledge certain trade-offs in our computational approach. To prevent bias and ensure broad accessibility, we prioritized unsupervised threshold-based segmentation over machine learning models that require labor-intensive manual labeling. Consequently, low-quality images—such as those with shading artifacts near well edges—are excluded rather than recovered, a limitation we mitigate through high experimental throughput. However, the annotated datasets generated by *Organoid Profiler*, containing thousands of verified masks, can serve as ground truth for training future deep-learning models. Meanwhile, the multi-feature metrics extracted can serve to provide large language models with explainable training data associated with organoid images. Additionally, while the pipeline detects distinct developmental phases, integrating real-time feedback loops to automatically trigger media changes and tune media components remains a future engineering goal.^19^

## Conclusion

We present a unified framework for the automated fabrication and quantitative profiling of organoids. By coupling the physical precision of microfluidics with the accessible power of *Organoid Profiler*, we have established a robust methodology that transforms organoid characterization from a qualitative art into a quantitative science. This platform not only improves the reproducibility of biological and biomedical research but also provides the standardized, high-content data streams necessary for drug discovery and personalized medicine.

## Methods

### Organoid Profiler

The image analysis workflow consists of three primary modules:

1. **Pre-processing and Segmentation:** Raw images (brightfield or fluorescence) are first converted to 8-bit depth. To address illumination inconsistencies and high-frequency noise, a Gaussian blur filter is applied. Images are binarized using automated thresholding algorithms (specifically the “Triangle” or “Otsu” methods, depending on contrast), followed by morphological operations including hole-filling, dilation, and erosion to define solid object boundaries. A watershed segmentation algorithm is applied to separate physically touching organoids.
2. **Artifact Exclusion:** To ensure data integrity, Regions of Interest (ROIs) are filtered based on user-defined or default criteria. Objects falling outside specific area ranges or circularity thresholds (0.0–1.0) are automatically excluded. Furthermore, ROIs located at the image periphery are removed to eliminate shadow artifacts and incomplete objects.
3. **Feature Extraction and Analysis:** For validated ROIs, the pipeline computes morphological metrics including Area, Perimeter, Circularity, and Solidity. For fluorescence images, Corrected Total Fluorescence (CTF) is calculated as *CTF* = Integrated Density of Organoid Regions − (Area of Organoids × Mean Fluorescence of Background). Data are exported as CSV files for downstream statistical analysis and longitudinal visualization.

*Organoid Profiler* source code and detailed description is available on GitHub (https://github.com/edgar-galan/organoid-profiler), and a web-based interface is accessible at https://www.organoid-profiler.com.

#### *Organoid Profiler* validation and comparison against manual measurements

To evaluate the accuracy and robustness of the *Organoid Profiler* segmentation algorithm, we established a validation dataset consisting of 240 representative organoid images selected from the longitudinal study. “Ground truth” measurements were generated by three independent researchers (H1, H2, and H3) who manually traced the boundaries of the same organoids using the “Polygon” selection tool in ImageJ (NIH). A “consensus” measurement was defined for each organoid as the arithmetic mean of the three manual annotations.

Statistical validation was performed using custom scripts written in Python (v3.12) utilizing the *SciPy* and *Pandas* libraries. Linearity between the automated *Organoid Profiler* measurements and the manual consensus was assessed using the Pearson correlation coefficient (*r*). Agreement between the methods was evaluated using Bland-Altman analysis, calculating the mean bias (average of the differences between automated and consensus measurements) and the 95% limits of agreement (LoA), defined as Bias ± 1.96 × s.d._diff_.

To benchmark the system’s precision against human performance, we quantified measurement variability. Inter-researcher variability was calculated for every pair of human annotators (H1 vs. H2, H1 vs. H3, H2 vs. H3) using the percentage difference metric:

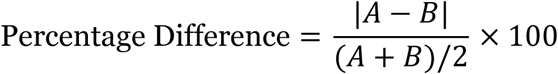

This was compared against the variability between the *Organoid Profiler* output and the human consensus, derived using the same metric. Distributions of these variability scores were visualized using violin plots to demonstrate that the automated system’s deviation from the consensus falls within the range of standard human-to-human variability.

#### Brightfield image processing and morphometric profiling

Brightfield images of organoids were processed using *Organoid Profiler*. The software automatically identifies organoid regions of interest and extracts a comprehensive set of morphological features, mathematically defined in **Extended Data Table 1**, including geometric parameters (e.g., Area, Perimeter, Aspect Ratio, Circularity, Roundness, Solidity), intensity-based metrics (e.g., Mean Intensity, Integrated Density), and distribution descriptors (e.g., Skewness, Kurtosis). Data is exported as CSV files for downstream statistical analysis.

#### Morphometric heatmaps

To visualize morphological trends over time, feature values were aggregated as the mean of the population for each time point. Clustered heatmaps were generated using the seaborn.clustermap function. Feature values were normalized row-wise using min-max scaling (0–1) to allow for comparison of relative trends across different units. Hierarchical clustering was applied to the features (rows) using Euclidean distance and average linkage, while time points (columns) were maintained in chronological order.

#### Time-course trajectory analysis

Longitudinal data were visualized as line plots representing the mean ± standard deviation (s.d.) superimposed over individual data points to display population distribution. For comparative analysis of variability, the Coefficient of Variation (CV, %) was calculated as (s.d./Mean) × 100 for all strictly positive morphological features (e.g., Area, Aspect Ratio). For features capable of taking values near zero (e.g., Skewness, Kurtosis), the standard deviation (s.d.) was used as the metric of variability. Intra-group variability was calculated specifically for each condition. To quantify the total variability of a heterogeneous population (“All conditions or batches”), data from all conditions were pooled, and statistics were computed across the combined dataset. Variability heatmaps were plotted using a diverging color palette to highlight stability versus heterogeneity.

#### Automated organoid segmentation and fluorescence quantification

Fluorescence microscopy images were processed using *Organoid Profiler* in fluorescence mode. Python OpenCV and SciPy libraries were used to extract morphological and fluorescence features from multi-channel image sets (Brightfield, Calcein-AM, and Propidium Iodide).

To define organoid boundaries, segmentation was performed on the fluorescence channels. Images were first denoised using a Gaussian blur (5 × 5 kernel). An initial binary mask was generated using Otsu’s automated thresholding method. To refine the segmentation masks and ensure the exclusion of debris and hollow centers, a sequence of morphological operations was applied: morphological opening (5 × 5 kernel), morphological closing (5 × 5 kernel), dilation (3 × 3 kernel), and erosion (3 × 3 kernel). Any remaining holes within the detected objects were filled using binary hole filling, and objects smaller than a set pixel threshold (*Valid* = *A* > *S_min_*; where *A* is Area of the ROI and *S_min_* is a minimum user-set size) were discarded.

Fluorescence intensity was quantified within the segmented organoid regions of interest (*R_roi_*) for both the Live and Dead channels. To account for local variations in background signal, a local background correction was applied to each image. The background region (*R_bg_*) was defined as the inverse of the organoid mask (all pixels not belonging to an organoid).

The mean background intensity (*μ_bg_*) was calculated as:

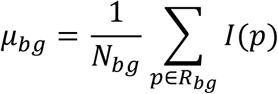

where *I*(*p*) represents the pixel intensity at position *p*, and *N_bg_* is the total number of pixels in the background region.

Two primary metrics were calculated for each organoid: *Corrected Mean Fluorescence* (*F_mean_*) and *Corrected Total Fluorescence* (*F_total_*).

*F_mean_* was derived by subtracting the background mean from the raw mean intensity of the ROI (*μ_roi_*):

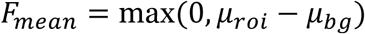

*F_total_* was calculated by subtracting the background contribution from the raw integrated density:

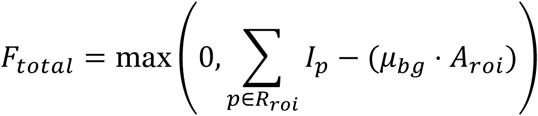

where *A_roi_* is the area of the organoid in pixels. These metrics (*F_mean_* and *F_total_*) served as the inputs for subsequent Live/Dead and Immunofluorescence analyses.

#### Live/Dead fluorescence quantification and statistical analysis

Quantitative fluorescence intensity data for Calcein-AM (live) and Propidium Iodide (PI; dead) were extracted using *Organoid Profiler* software. Data were analyzed using the *F_mean_* and *F_total_* metrics derived from the image processing pipeline.

To enable comparisons across datasets with varying baseline intensities, a global Z-score normalization was applied using a pooled dataset of all experimental conditions.

A global mean (*μ_blobal_*) and global standard deviation (*σ_global_*) were calculated from the pooled *F_mean_* values of both channels. The normalized fluorescence (*F_norm_*) for each individual organoid *j* was calculated as:

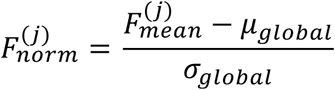

For visualization, normalized mean fluorescence data were plotted as individual data points (strip plots) overlayed with the daily mean ± standard deviation. To visualize the relationship between organoid size and fluorescence intensity, individual data points were color-coded based on their corrected total fluorescence intensity (low to high gradient). The analysis spanned 11 days for microfluidics-engineered organoids and selected time points (Days 4, 7, and 11) for non-engineered controls.

#### Immunofluorescence staining quantification and statistical analysis

Quantitative immunofluorescence data were processed using the *F_mean_* and *F_total_* metrics described above. Samples were categorized into “Microfluidics-Engineered Organoids”, “Non-Engineered Organoids”, or “Tissue” controls based on input file nomenclature. Native tissue samples served as the baseline reference (Day 0). To account for experimental variability and enable direct comparisons, fluorescence intensity for both microfluidics-engineered organoids and non-engineered organoids was normalized to the native tissue control for each specific marker.

To evaluate expression relative to native tissue, *F_mean_* was normalized to the tissue control group for each specific marker. The mean fluorescence intensity of the tissue controls (*μ_tissue_*) was calculated as:

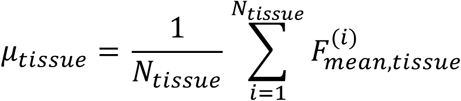

where *N_tissue_* is the number of tissue ROIs measured. The relative normalized fluorescence (*F_norm_*) for every sample *j* was then determined as the fold-change over tissue:

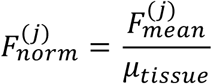

Data were visualized on a common shared y axis for both microfluidics-engineered organoids and non-engineered organoids post-normalization, enabling direct comparison. Individual data points were displayed as strip plots, where the y-axis represented the normalized mean fluorescence 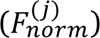 To provide additional dimensionality, individual points were color-coded based on their *Corrected Total Fluorescence* 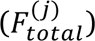. Native tissue samples were assigned to Day 0 to serve as the baseline reference and were plotted as discrete points, while organoid developmental trajectories (Day > 0) were visualized as connected lines. Error bars indicate the mean ± standard deviation (s.d.). A reference line at *y* = 1.0 was included to indicate the baseline expression level of the native tissue.

### Animal and tissue sourcing

Mice were housed in a pathogen-free environment at 23 ± 2 °C and 50–65% relative humidity under a 12/12 h light/dark cycle with free access to food and water, in accordance with the National Institutes of Health guidelines. Lung and liver organoids were derived from 6–8-week-old male BALB/c mice. Mice were euthanized via CO₂ asphyxiation in accordance with ethical guidelines of the Animal Experimentation Ethics Committee at Tsinghua Shenzhen International Graduate School (SIGS), Tsinghua University (Project No: 2022/123, 2024/F021). Lung and liver tissues were surgically excised, minced into approximately 1–3 mm^3^ fragments, and rinsed three times in ice-cold phosphate-buffered saline (PBS). To prevent contamination, tissue samples were treated with 2% (v/v) penicillin-streptomycin (Gibco). Tissue digestion was performed using a cocktail of collagenase I (2 mg mL^−1^; Sigma Aldrich), collagenase IV (2 mg mL^−1^; Sigma Aldrich), dispase II (2 mg mL^−1^; Sigma Aldrich), and DNase I (200 U mL^−1^; Sigma Aldrich) supplemented with 1% penicillin-streptomycin in Dulbecco’s Modified Eagle Medium/Nutrient Mixture F-12 (DMEM/F12; Gibco). The resulting cell suspension was filtered through a 100 *μ*m cell strainer (Falcon) to remove undigested debris. Digestion was halted by resuspending the filtrate in DMEM supplemented with 10% fetal bovine serum (FBS). Cells were centrifuged at 300 × *g* for 5 minutes, and the supernatant was discarded. Red blood cell lysis was performed using a specific lysis buffer (1 mL per 1 × 10^6^ cells) with an incubation time of 3 minutes. Following a final centrifugation at 300 × *g* for 5 minutes, cell viability and concentration were assessed using the Trypan blue exclusion method (Gibco).

### Human cerebral organoid generation

Brain organoids were generated from human pluripotent stem cells (hPSCs) following the protocol described by Lancaster and Knoblich.^20^ Briefly, hPSCs were dissociated into single cells and plated in low-attachment 96-well plates to facilitate the formation of embryoid bodies (EBs). On Day 6, EBs were transferred to neural induction media to promote neuroectodermal differentiation. On Day 11, neuroepithelial tissues were embedded in droplets of Matrigel (Corning) and transferred to a spinning bioreactor to enhance nutrient absorption and tissue expansion. Organoids were maintained in differentiation media at 37°C with 5% CO₂ until imaging.

### Microfluidic organoid bioprinting and culture

Automated organoid fabrication was performed using the OrgFab-P01 system^21^ (SynOrg Biotechnology; Shenzhen, China). Single cells derived from mouse tissue or hPSCs were suspended in Matrigel at optimized concentrations. The cell-laden hydrogel suspension was loaded into the microfluidic printing system and co-flowed with HFE-7000 oil (fluorinated oil) into a three-way flow-focusing microfluidic device. This geometry facilitated the generation of monodisperse droplets, which were transported through polytetrafluoroethylene (PTFE) tubing. The tubing passed through a temperature-controlled stage maintained at 37 °C to induce Matrigel polymerization (gelation). The solidified organoid droplets were subsequently ejected through a nozzle into 96-well culture plates. Organoids were incubated at 37 °C with 5% CO₂.

### Manual organoid fabrication

For conventional dome cultures, the cell-Matrigel suspension was manually pipetted as 20–50 μl droplets directly onto the surface of 6-well culture plates. Plates were inverted to prevent cell settling and incubated at 37 °C for 15–20 minutes to allow gelation before the addition of culture medium.

### Live/Dead fluorescence staining

To assess organoid viability and morphology, live and dead cell staining was performed using Calcein-AM and Propidium Iodide (PI) (Thermo Fisher Scientific). Organoids were washed three times with PBS to remove residual culture medium. A staining solution was prepared by diluting Calcein-AM (final concentration 2 *μ*M) and PI (final concentration 1.5 *μ*M) in PBS. 50 *μ*L of the staining solution was added to each well of a 96-well plate containing a single organoid. Samples were incubated at 37 °C for 25 min protected from light. Imaging was performed immediately using a Nikon Eclipse Ts2r inverted fluorescence microscope. Fluorescence images were captured at two focal planes to encompass both the organoid outline and core; average fluorescence intensities were calculated from these planes.

### Luciferase viability assay

Longitudinal metabolic viability was quantified using the CellTiter-Glo Luminescent Cell Viability Assay (Promega), following the manufacturer’s instructions. At each indicated time point, organoids were incubated with the CellTiter-Glo reagent (1:1 volume ratio) to lyse cells and generate a luminescent signal proportional to the amount of ATP present. Luminescence was measured using a microplate reader. The assay was performed as an endpoint measurement on independent batches (*n* = 3 biological replicates) for each time point. Data processing was performed using custom Python scripts: raw Relative Light Units (RLU) were first background-corrected by subtracting the mean signal of blank wells, then normalized to the Day 0 baseline of the corresponding biological replicate to determine the fold-change in viability over time.

### Histological analysis (H&E staining)

For morphological assessment, organoids were fixed in 4% paraformaldehyde (PFA) at room temperature for 2 h, rinsed in PBS, and processed for cryosectioning. Cryosections (8 *μ*m thickness) were prepared and mounted on glass slides. Sections were dehydrated through a graded ethanol series (70%, 80%, 90%, and 100%) and stained with eosin for 2 min to visualize the cytoplasm and extracellular matrix. Slides were rinsed in water and treated with a bluing solution to enhance nuclear contrast. Following staining, sections were dehydrated, cleared in 100% xylene, and mounted with an antifade mounting medium. Brightfield images were acquired using a high-resolution light microscope.

### Flow cytometry analysis

Organoids were harvested at Days 4, 7, 9, and 11 and dissociated into single-cell suspensions using TrypLE Express (Gibco) for 15 min at 37 °C with mechanical disruption. Native lung tissue was digested using the same protocol to establish a Day 0 baseline. Cells were washed with FACS buffer (PBS with 2% FBS and 2 mM EDTA) and blocked with FcR blocking reagent (Miltenyi Biotec) for 10 min at 4 °C. Staining was performed using the following fluorophore-conjugated primary antibodies (BioLegend): anti-CD45 (Leukocytes), anti-CD31 (Endothelial), anti-CD326/EpCAM (Epithelial), anti-MHC Class II (I-A/I-E), anti-F4/80, and anti-CD11b. Viability was assessed using 7-AAD.

Data were acquired on a CytoFLEX LX flow cytometer (Beckman Coulter) and analyzed using FlowJo software (v10.8). The gating strategy was defined as follows: Debris and doublets were excluded using FSC/SSC parameters. Viable cells (7-AAD^−^) were stratified into leukocytes (SSC-A low/med CD45^+^) and somatic cells (CD45^−^). Within the CD45^−^ fraction, endothelial cells were identified as CD31^+^ CD326^−^ and epithelial cells as CD31^−^ CD326^+^. The epithelial population was further stratified into alveolar cells (MHC II^+^ CD326^+^) and airway cells (MHC II^−^ CD326^+^). Macrophages were identified within the CD45^+^ fraction as F4/80^+^ CD11b^+^. See **Supplementary Table 3** for the list of fluorophore-conjugated antibodies used for flow cytometry.

### Quantitative real-time PCR (qPCR)

Total RNA was extracted from organoids and native lung tissue using the RNeasy Mini Kit (Qiagen) and reverse-transcribed using the iScript cDNA Synthesis Kit (Bio-Rad). qPCR was performed on a CFX96 Touch Real-Time PCR Detection System (Bio-Rad) using iTaq Universal SYBR Green Supermix (Bio-Rad). Relative gene expression was determined using the comparative *C_T_* method (ΔΔ*C_T_*) normalized to Gapdh and calibrated to native mouse lung tissue (Day 0) for the lineage markers: *Nkx2-1*, *Krt5*, *Krt15*, *Muc5a*, *Scgb1a1*, *Lamp3*, *Sftpc*, *Ager*, *Cav1*, *Rtkn2*. Statistical significance was determined using one-way ANOVA with Tukey’s multiple comparison test. See **Supplementary Table 4** for the list of oligonucleotide primers used for qPCR.

### Immunofluorescence staining

Organoids were fixed in 4% paraformaldehyde (PFA) for 20 min, permeabilized with 0.5% Triton X-100/PBS for 15 min, and blocked with 5% donkey serum for 1 h. Samples were incubated overnight at 4 °C with primary antibodies against Nkx2-1 (1:100, Abcam), Sftpc (1:200, Millipore), Hopx (1:200, Santa Cruz), Cd68 (1:100, Abcam), Cd3 (1:100, BioLegend), and *α*-SMA (1:200, Sigma). Detection was performed using Alexa Fluor 488-, 568-, or 647-conjugated secondary antibodies (1:500, Thermo Fisher Scientific). Nuclei were counterstained with DAPI. Images were acquired using a Leica SP8 confocal microscope. See **Supplementary Table 2** for the list of primary and secondary antibodies used for immunofluorescence staining.

### RNA-sequencing

#### RNA extraction and library preparation

Total RNA was extracted from microfluidic organoids collected at Days 1, 4, 7, and 11, and from native mouse lung tissue using TRIzol reagent (Invitrogen) with standard protocols. RNA integrity and concentration were assessed prior to library preparation. Sequencing libraries were constructed using the Illumina TruSeq Stranded mRNA Library Prep Kit according to the manufacturer’s instructions.

#### RNA sequencing and upstream processing

Sequencing was performed on an Illumina platform (Novogene). Raw reads were processed to remove adapter sequences and low-quality bases. Clean reads were aligned to the mouse reference genome using HISAT2. Gene expression levels were quantified by counting reads mapped to each gene using the featureCounts tool from the Subread package.

#### Differential gene expression analysis

Differential expression analysis between organoids and native tissue was performed using the DESeq2 R package. Raw counts were normalized using the median of ratios method. Log2 fold changes for time-course samples (Day 1, 4, 7, 11) were calculated relative to native lung tissue. Principal Component Analysis (PCA) was performed on log2-transformed TPM values using scikit-learn. Gene Set Enrichment Analysis (GSEA) was conducted using gseapy with custom gene sets for “Immune response,” “Lung identity,” and “Matrix remodeling.” Enriched terms were considered significant at FDR *q* < 0.05. Heatmaps were generated using seaborn with hierarchical clustering (Euclidean distance, Ward’s linkage). Genes with an adjusted *P* value < 0.05 and an absolute log_2_ fold change > 1 were considered significantly differentially expressed. Volcano plots were generated using Python (Matplotlib/Seaborn) to visualize the global gene expression changes, highlighting key upregulated and downregulated genes.

#### Functional enrichment analysis

To characterize the biological functions associated with the differentially expressed genes, functional enrichment analysis was performed for Kyoto Encyclopedia of Genes and Genomes (KEGG) pathways and Gene Ontology (GO) biological process terms.

#### KEGG analysis

Enrichment of KEGG pathways was assessed using the GSEApy library (v1.0.4) against the “KEGG_2019_Mouse” gene set database. Pathways with an adjusted *P* value < 0.05 were considered significant. The top enriched pathways were visualized as a dot plot, where dot size represents the number of differentially expressed genes (count) and color indicates statistical significance (adjusted *P* value).

#### GO analysis

Gene Ontology enrichment for biological processes was performed using GSEApy against the “GO_Biological_Process_2023” database. To reduce redundancy and improve plot readability, a curated list of representative terms was selected for visualization. Enriched terms were filtered based on an adjusted *P* value < 0.05. Results were visualized as dot plots and bar plots to illustrate the temporal dynamics of processes related to proliferation, differentiation, and tissue organization.

#### Statistical analysis for RNA-sequencing

For all RNA-seq data, *P* values were adjusted for multiple testing using the Benjamini-Hochberg procedure. All plotting and downstream data processing were performed using custom Python scripts utilizing Pandas (v1.3.5), Matplotlib (v3.5.1), and Seaborn (v0.11.2).

## Ethics statement

All animal experiments were conducted in compliance with relevant ethical regulations regarding the use of research animals and were approved by the Animal Experimentation Ethics Committee at Tsinghua Shenzhen International Graduate School (SIGS), Tsinghua University (Project No: 2022/123, 2024/F021). Male BALB/c mice were acquired from the Guangdong Medical Laboratory Animal Center and housed at the Peking University Laboratory Animal Center, University Town of Shenzhen.

## Competing interests

Shaohua Ma and Yongde Cai are co-founders of SynOrg Biotechnology (Shenzhen).

## Data availability

All datasets associated with this publication are available at Zenodo; currently, a list of Zenodo links are found at (https://github.com/edgar-galan/organoid-profiler). The source data underlying all Figures and Extended Data Figures are provided as a Source Data file with this paper. Requests for materials should be addressed to.

## Code availability

*Organoid Profiler* source code is available on GitHub (https://github.com/edgar-galan/organoid-profiler), and a web-based interface is accessible at https://www.organoid-profiler.com. Python scripts used to process, analyze and visualize the data are currently available at https://github.com/edgar-galan/organoid-profiler. Technical queries can be addressed directly to.

## Acknowledgements

The authors thank the National Key Research and Development Program of China (2024YFA0919800), Shenzhen Medical Academy of Research and Translation (SMART; B2402009), National Natural Science Foundation of China (32371470, 82341019), Shenzhen Natural Science Foundation (JCYJ20241202123909013), and the Key-Area Research and Development Program of Guangdong (2023B0909020003).

## Author Contributions

E.A.G.: Conceptualization, Software, Investigation, Formal analysis, Visualization, Writing – original draft, Writing – review & editing, Methodology, Validation, Data Curation. W.W.: Investigation. Y.Z.: Investigation. Z.W.: Investigation. J.W.: Investigation. A.N.F.: Software. Y.C.: Resources. G.S.: Investigation. X.D.: Resources. S.M.: Conceptualization, Supervision, Funding acquisition, Writing – review & editing.

## Supplementary Information

Supplementary Information is available for this paper.

## Extended Data

**Extended Data Table 1:**
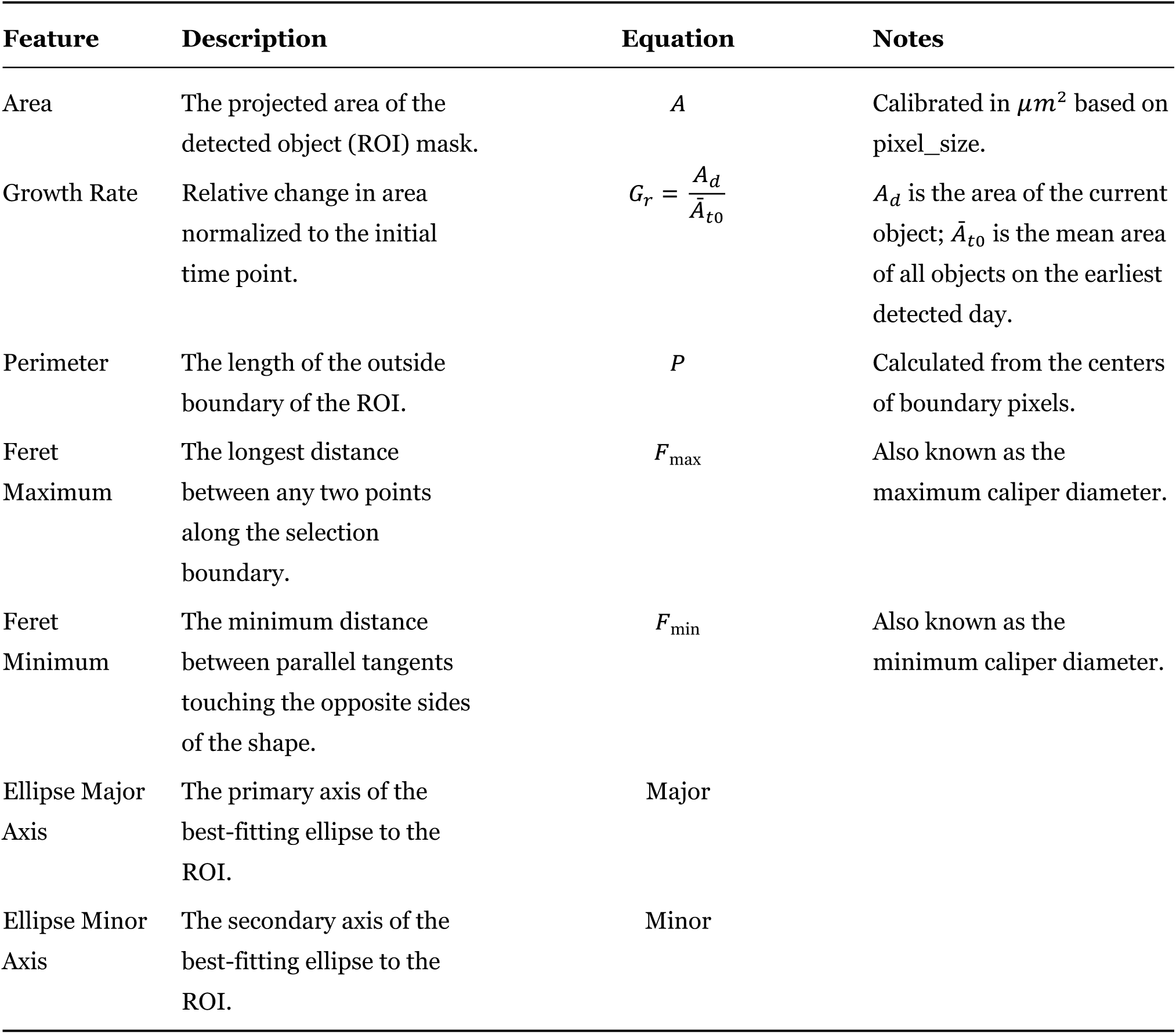
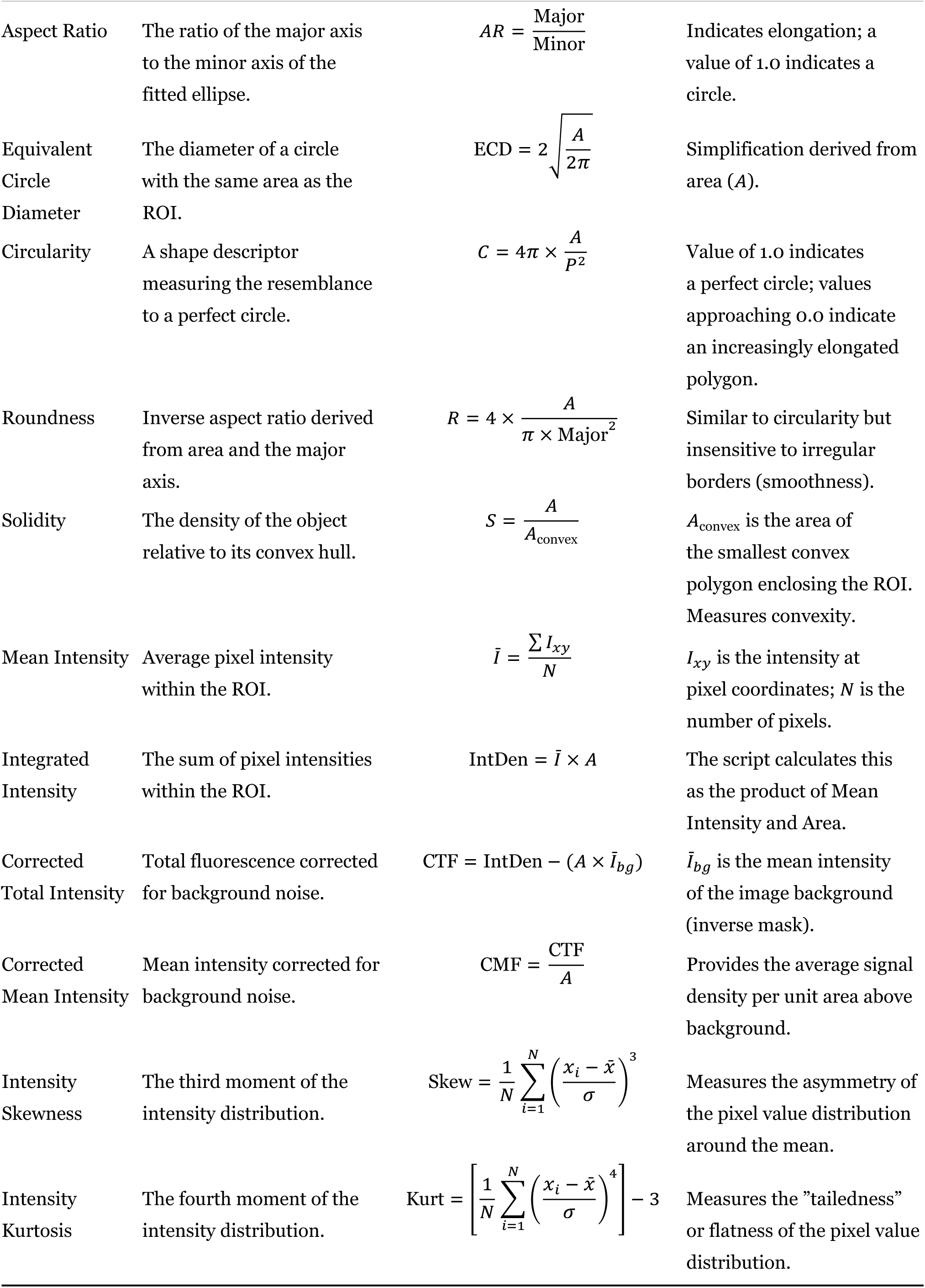
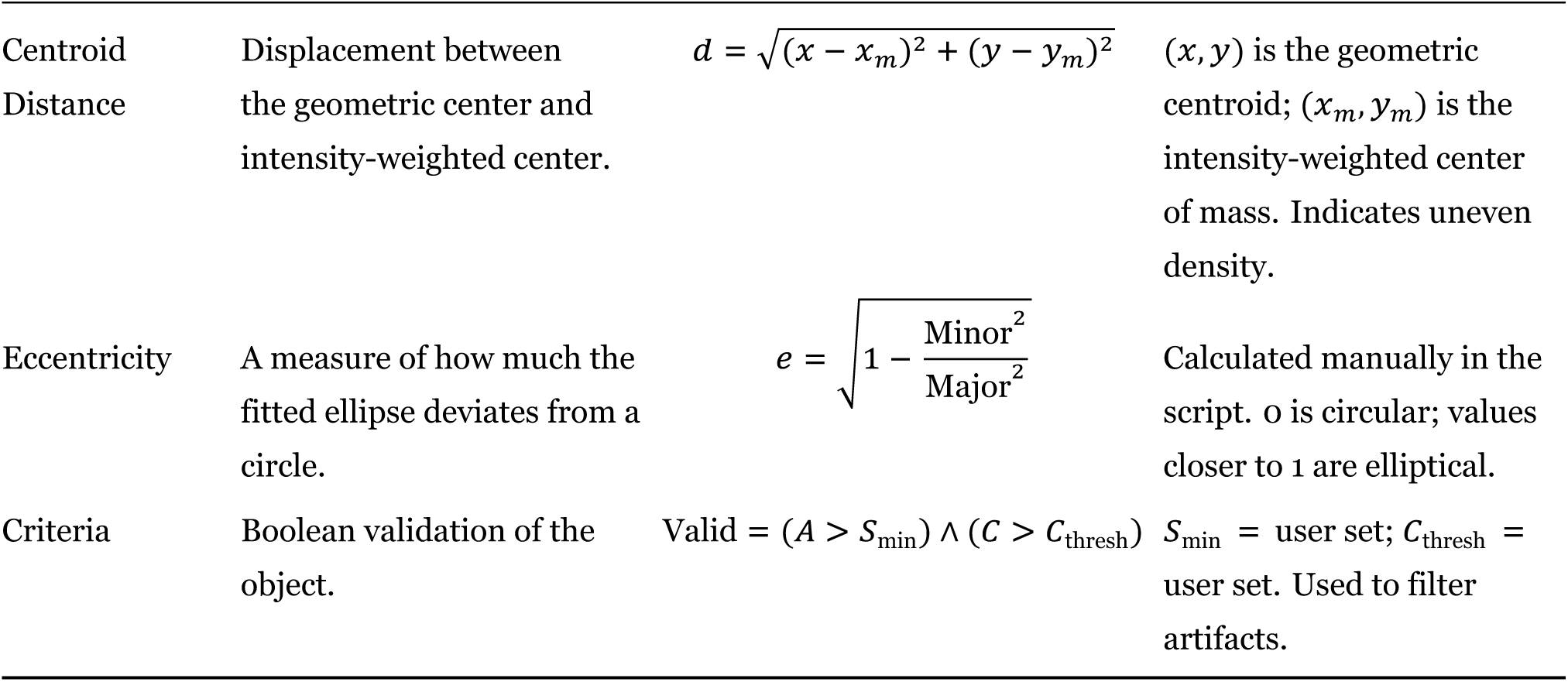
Mathematical definitions and descriptions of morphometric features extracted by *Organoid Profiler*. Comprehensive list of the quantitative parameters automatically calculated by the image analysis pipeline to characterize organoid geometry and optical density. Columns include the Feature name, a biological or physical Description, the mathematical Equation used for computation, and technical Notes regarding implementation or units. Geometric features (e.g., Area, Circularity, Solidity) are derived from the binary mask of the segmented region of interest, while intensity-based metrics (e.g., Corrected Total Intensity, Skewness, Kurtosis) are calculated from pixel intensity values within the region of interest applied to the original 8-bit image. Abbreviations: ROI, region of interest; *A*, area; *P*, perimeter; *I*, intensity; *N*, number of pixels; s.d., standard deviation.

**Extended Data Fig. 1:**
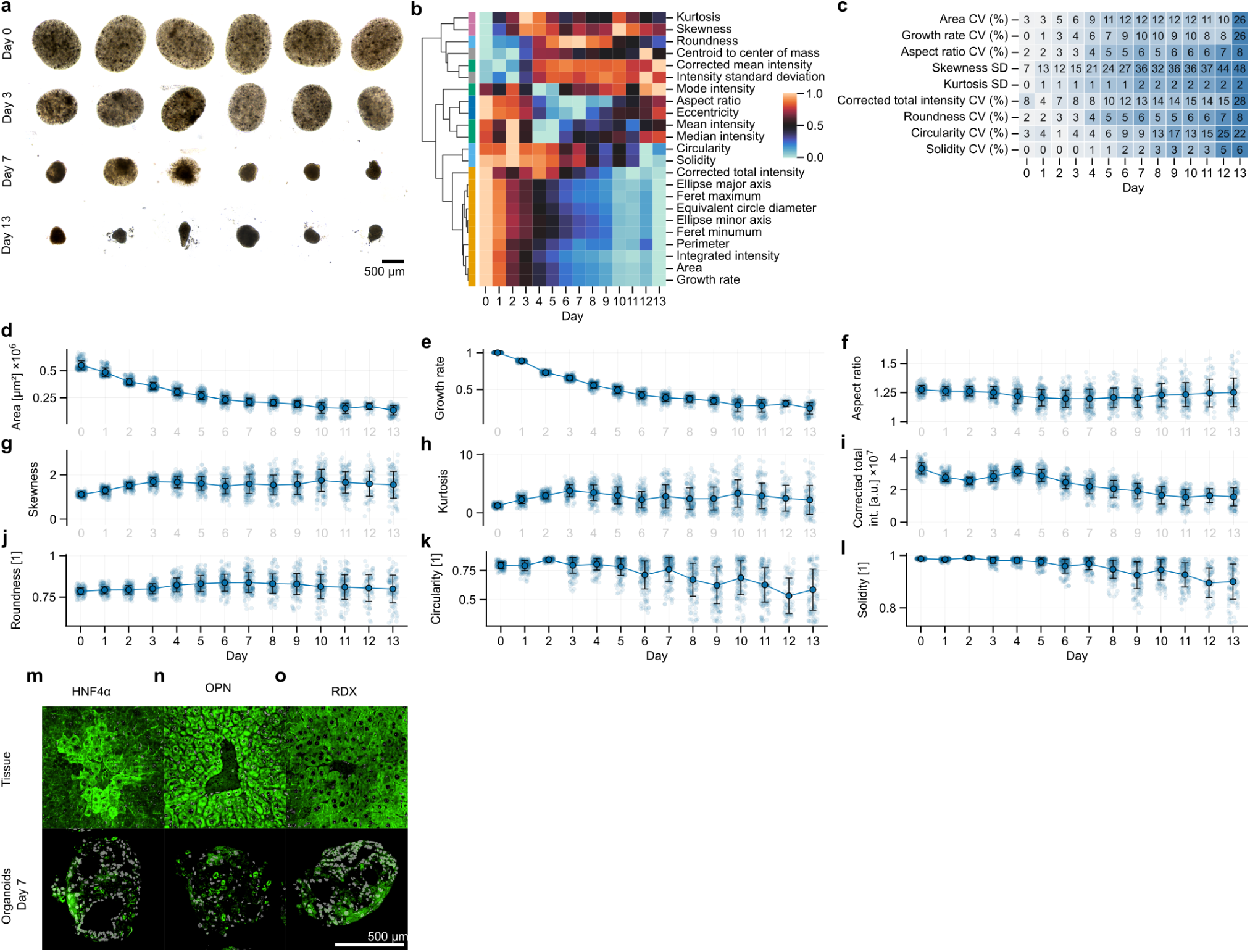
Morphometric profiling of microfluidics-engineered primary mouse liver organoids. **a** Representative brightfield images of the same six organoids tracked over time at Days 0, 3, 7, and 13 of culture. Scale bar, 500 *μ*m. **b** Clustered heatmap showing the morphometric profile of organoids (*n* = 192 biologically independent samples). **c** Heatmaps showing the coefficient of variation (CV) or standard deviation (s.d.) for Area, Growth rate, Aspect ratio, Skewness, Kurtosis, Corrected total intensity, Roundness, Circularity, and Solidity. **d**–**l** Line plots with overlapped strip plots showing the temporal evolution of **d** Area, **e** Growth rate, **f** Aspect ratio, **g** Skewness, **h** Kurtosis, **i** Corrected total intensity, **j** Roundness, **k** Circularity, and **l** Solidity over 13 days. Data are presented as mean values; individual data points represent biologically independent organoids (*n* = 96). **m–o** Representative immunofluorescence images of mouse liver tissue and primary mouse liver organoids at Day 7 stained for **m** the hepatocyte marker HNF4α, **n** the cholangiocyte marker OPN, and **o** the apical marker RDX. Scale bar, 500 *μ*m. Source data are provided as a Source Data file.

**Extended Data Fig. 2:**
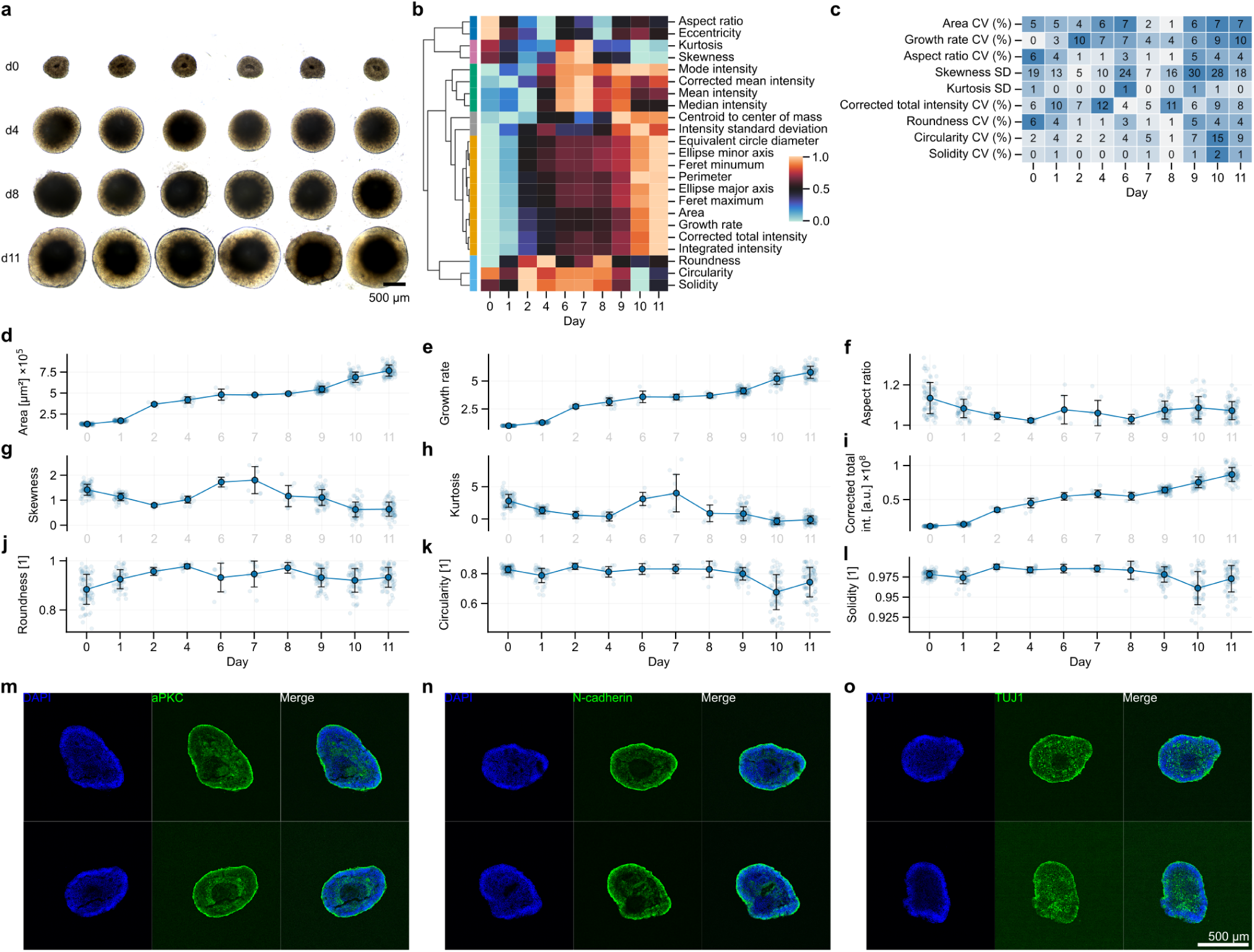
Evaluation of organoid morphology and batch variation. **a** Representative brightfield images of a single batch of hiPSC-derived cerebral organoids at Days 0, 4, 8, and 11 of differentiation. The same six organoids were tracked over time. Scale bar, 500 *μ*m. **b** Clustered heatmap (“morphometric profile”) of morphological features extracted from brightfield images (*n* = 96 organoids). **c** Heatmap showing the coefficient of variation (CV) or standard deviation (s.d.) of selected morphological features (Area, Growth rate, Aspect ratio, Skewness, Kurtosis, Corrected total intensity, Roundness, Circularity, and Solidity) across differentiation days. **d–l** Temporal quantification of morphological features: **d** Area, **e** Growth rate, **f** Aspect ratio, **g** Skewness, **h** Kurtosis, **i** Corrected total intensity, **j** Roundness, **k** Circularity, and **l** Solidity. Data are presented as mean ± s.d. with individual data points overlaid (*n* = 96 organoids). **m–o** Representative immunofluorescence images of cerebral organoids at Day 11 stained for **m** the apical marker aPKC, **n** the neural progenitor marker N-cadherin, and **o** the neuronal marker TUJ1. Scale bar, 500 *μ*m. Source data are provided as a Source Data file.

**Extended Data Fig. 3:**
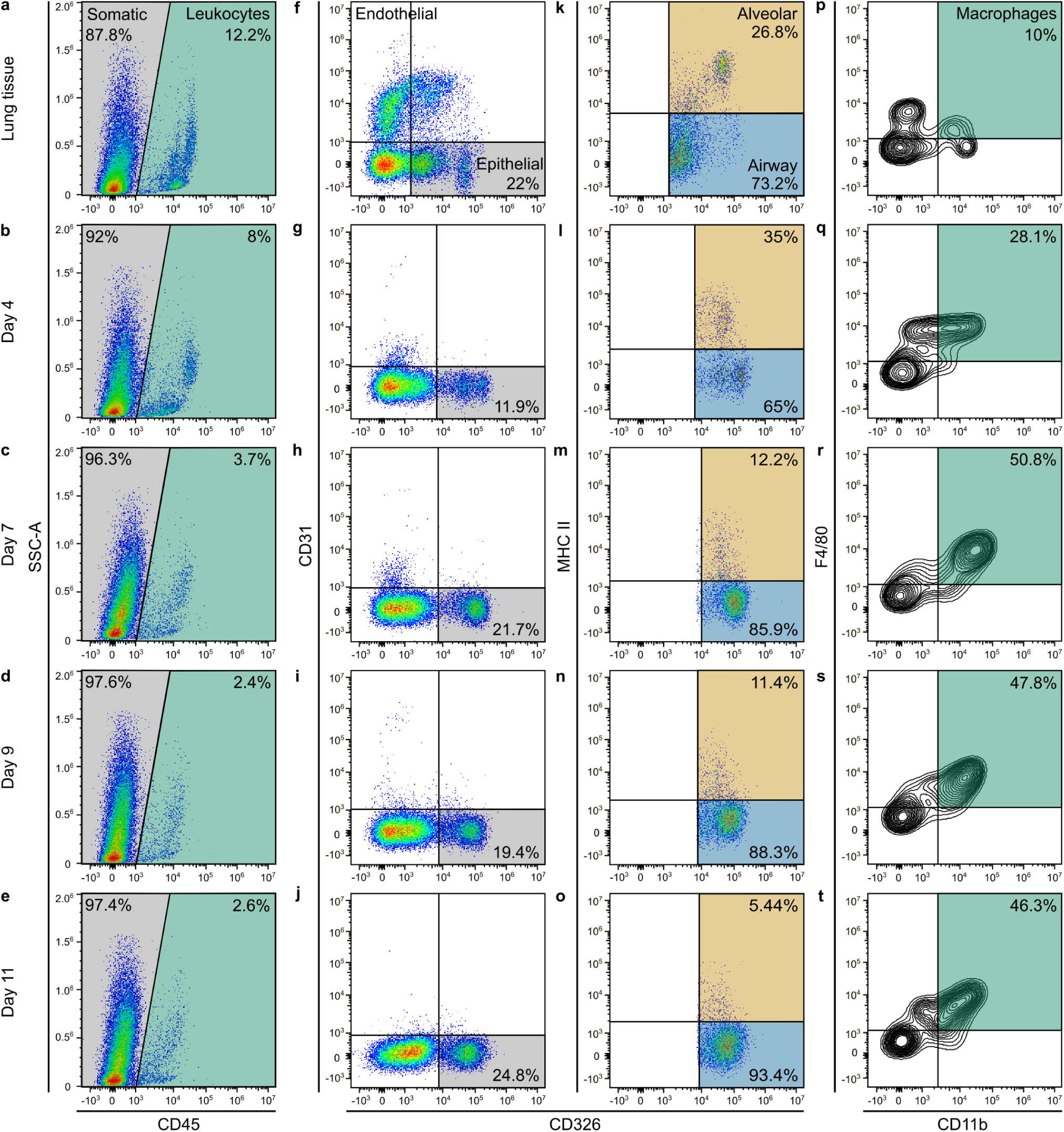
Flow cytometry gating strategy for quantification of cell populations in microfluidics-engineered organoids and native lung tissue. Representative flow cytometry density plots illustrating the hierarchical gating logic used to identify distinct cell lineages in digested native mouse lung tissue (top row) and microfluidics-engineered organoids during differentiation at Days 4, 7, 9, and 11 (rows 2–5). **a–e** Identification of Somatic (CD45^−^) and Leukocyte (CD45^+^) populations based on CD45 expression and side scatter (SSC-A). **f–j** Discrimination of Endothelial (CD45^−^CD326^−^CD31^+^) and Epithelial (CD45^−^CD31^−^CD326^+^) cells within the somatic gate. **k–o** Further stratification of the CD326^+^ epithelial population into Alveolar (MHCII^+^) and Airway (MHCII^−^) lineages. **p–t** Identification of Macrophages (CD45^+^CD11b^+^F4/80^+^) within the leukocyte gate. Numbers within the plots indicate the percentage of cells in the respective gates relative to the parent population. This gating strategy was used to generate the quantitative data presented in **Fig. 6d–h**.

**Extended Data Fig. 4:**
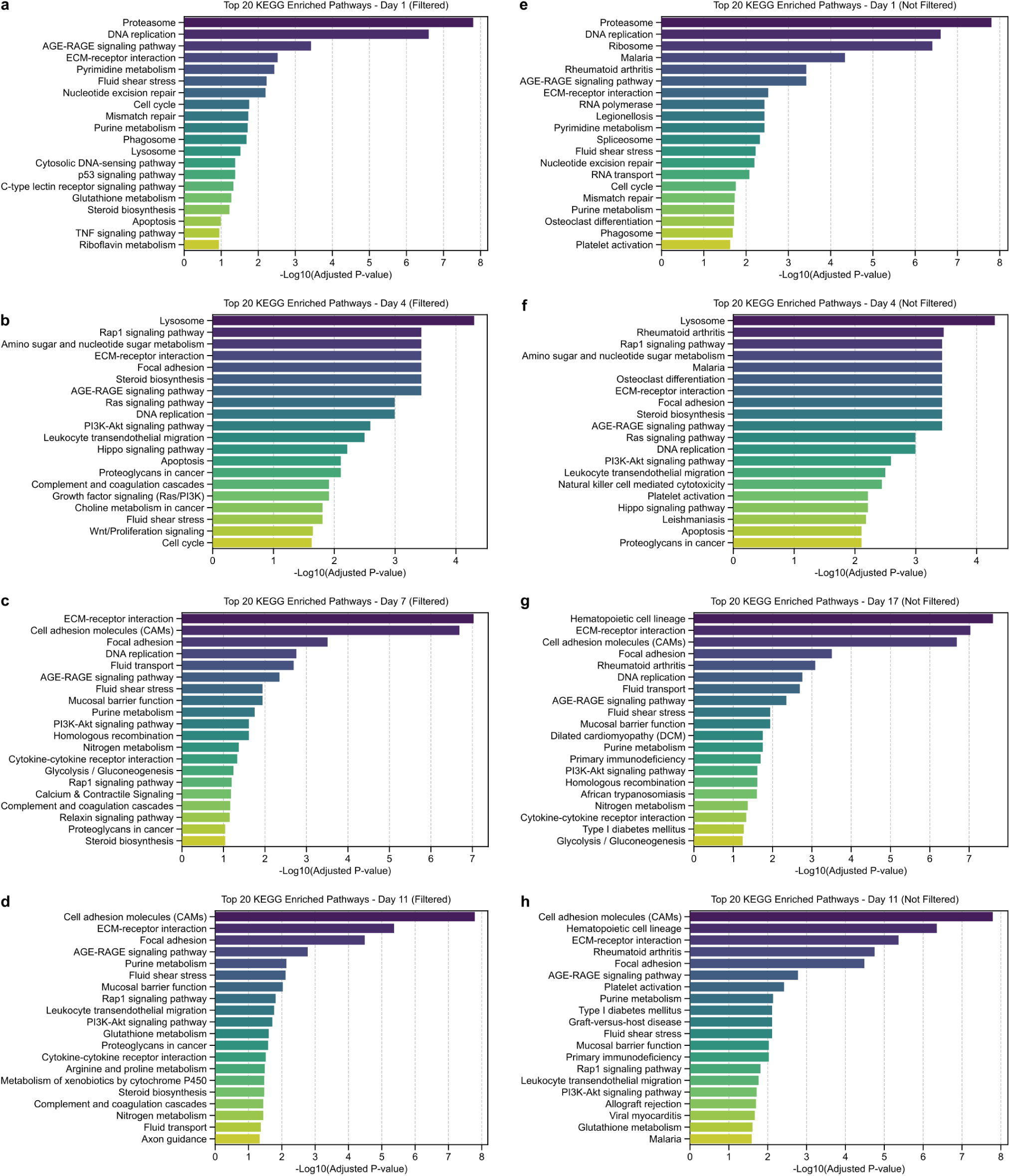
Comparison of filtered and unfiltered KEGG pathway enrichment analysis. **a–h** Bar plots displaying the top 20 significantly enriched KEGG pathways in microfluidics-engineered primary mouse lung organoids compared to native mouse lung tissue, ranked by statistical significance (− log_10_(adjusted *P* value)). Panels **a**–**d** show the results after filtering to remove contextually irrelevant or redundant terms (e.g., specific infectious disease pathways not relevant to the model) for **a** Day 1, **b** Day 4, **c** Day 7, and **d** Day 11. Panels **e**–**h** show the corresponding unfiltered results for **e** Day 1, **f** Day 4, **g** Day 7, and **h** Day 11, presenting the raw top 20 enriched pathways. The filtering step highlights biological processes specific to tissue development and maturation (e.g., ECM-receptor interaction, cell adhesion) by removing broad or non-specific entries present in the unfiltered lists. Source data are provided as a Source Data file.

**Extended Data Fig. 5:**
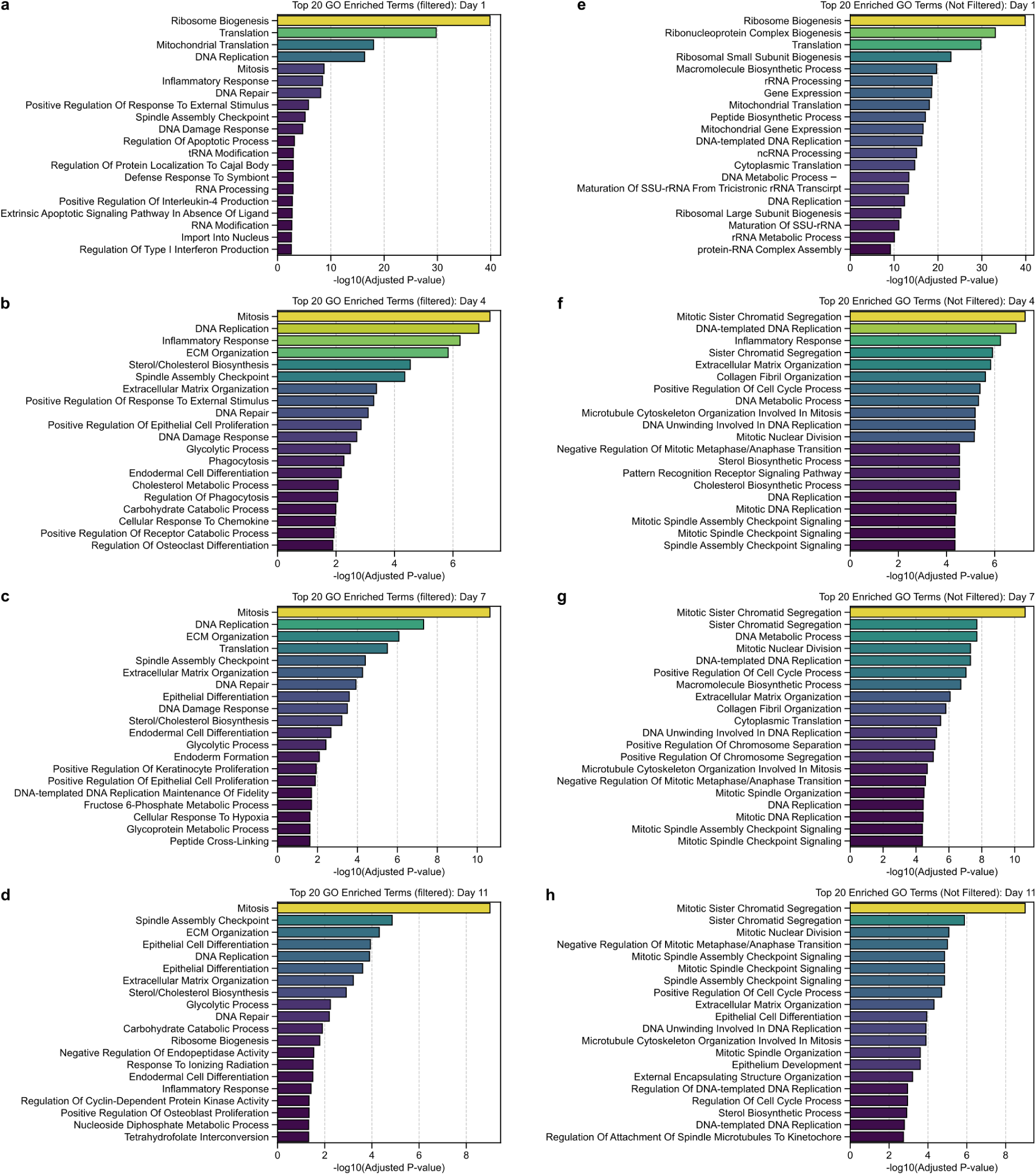
Comparison of filtered and unfiltered Gene Ontology (GO) biological process enrichment analysis. **a–h** Bar plots displaying the top 20 significantly enriched GO biological process terms in microfluidics-engineered primary mouse lung organoids compared to native mouse lung tissue, ranked by statistical significance (− log_10_(adjusted *P* value)). Panels **a**–**d** show the results after filtering to remove redundant or broad terms (e.g., general metabolic processes) and consolidating related biological functions for **a** Day 1, **b** Day 4, **c** Day 7, and **d** Day 11. Panels **e**–**h** show the corresponding unfiltered results for **e** Day 1, **f** Day 4, **g** Day 7, and **h** Day 11, presenting the raw top 20 enriched terms. The filtering process highlights specific developmental and structural programs (e.g., epithelial differentiation, tissue morphogenesis) relevant to the organoid model that might otherwise be obscured by highly significant but generic housekeeping processes in the unfiltered lists. Source data are provided as a Source Data file.

**Extended Data Video 1: Spontaneous contractions of alveolar structures driven by skeletal muscle lineage plasticity.** Time-lapse microscopy of microfluidics-engineered organoids (Day 7). Spontaneous, rhythmic contractions were observed in alveolar structures. RNA-seq analysis of these organoids revealed a specific upregulation of embryonic skeletal muscle contractile machinery (*Myh3*, *Myh8*, *Tnni1*) and a downregulation of airway smooth muscle markers (*Acta2*, *Myh11*) relative to lung tissue. This suggests the contractions are driven by a subpopulation of progenitors undergoing *de novo* striated muscle differentiation, reflecting the developmental multipotency of the foregut mesenchyme, rather than myofibroblast or cardiac activity.

## Notes

https://github.com/Edgar-Galan/organoid-profiler/

https://www.organoid-profiler.com/

