## Supplementary Information for "Organoid Profiler: Automated, high-throughput and quantitative morphological characterization uncovers conserved longitudinal developmental kinetics in microfluidics-engineered organoids"

**Supplementary Table 1. Composition of the custom primary mouse lung organoid culture medium.**

| Reagent | Final Concentration |
|---|---|
| DMEM/F12 | 86% (v/v) |
| Fetal Bovine Serum (FBS) | 10% (v/v) |
| B27 Supplement | 2% (v/v) |
| GlutaMax | 1% (v/v) |
| Penicillin-Streptomycin (P/S) | 1% (v/v) |
| Noggin | 100 ng $mL^{-1}$ |
| R-spondin 1 | 100 ng $mL^{-1}$ |
| Y27632 | 5 $\mu$M |
| CHIR99021 | 5 $\mu$M |

| Reagent | Final Concentration |
|---|---|
| EGF | 20 ng $mL^{-1}$ |
| FGF4 | 10 ng $mL^{-1}$ |
| cAMP | 20 $\mu$M |
| SB431542 | 10 $\mu$M |

### Supplementary Table 2. Primary and secondary antibodies used for immunofluorescence staining

| Target Antigen | Manufacturer | Catalog Number |
|---|---|---|
| F4/80 | Sigma-Aldrich | SAB5500103 |
| EpCAM | Sigma-Aldrich | HPA026761 |
| SFTPC | Sigma-Aldrich | 665984 |
| MUC5AC | Sigma-Aldrich | WH0004586M7 |
| $\alpha$-SMA | Sigma-Aldrich | F3777 |
| TUBB3 | Sigma-Aldrich | SAB4700544 |

### Supplementary Table 3. Fluorophore-conjugated antibodies used for flow cytometry

| Antibody / Reagent | Manufacturer | Catalog Number |
|---|---|---|
| 7-AAD Viability Staining Solution | BioLegend | 420404 |
| FITC anti-mouse CD45 | BioLegend | 103108 |
| Brilliant Violet 510™ anti-mouse F4/80 | BioLegend | 123135 |
| APC/Fire™ 750 anti-mouse CD31 | BioLegend | 102434 |
| Brilliant Violet 510™ anti-mouse CD90.2 | BioLegend | 140319 |
| PE/Cyanine7 anti-mouse I-A/I-E | BioLegend | 107630 |
| Brilliant Violet 421™ anti-mouse CD326 (EpCAM) | BioLegend | 118225 |
| APC/Fire™ 750 anti-mouse/human CD11b | BioLegend | 101262 |

### Supplementary Table 4. Oligonucleotide primers used for RT-qPCR

| Gene | Forward Sequence (5’–3’) | Reverse Sequence (5’–3’) |
|---|---|---|
| *Sftpc* | GAGGTCCTGATGGAGAGTCC | CCACGATGAGAAGGCGTTTG |
| *Lamp3* | GTGGCACCCGAAAATCCAAC | TGTCTGGAACATCACCACCG |
| *Nkx2.1* | TGCCGTGTAAACACGAGGAC | GACTCTCAAGCAAGTCCATCC |

| Gene | Forward Sequence (5'–3') | Reverse Sequence (5'–3') |
|---|---|---|
| *Ager* | ACTACCGAGTCCGAGTCTACC | CCCACCTTATTAGGGACACTGG |
| *Pdpn* | ACCCCAATAGAGATGGCTTGC | TAGGGCGAGAACCTTCCAGA |
| *Cav1* | TGAGAAGCAAGTGTATGACGC | CTTCCAGATGCCGTCGAAAC |
| *Muc5ac* | TGCATGCGTACCTGCCAGAA | CACACTGCATTGTGCCCTCA |
| *Scgb1a1* | CAAAAGCCCAGAGAAAGCATC | CAGTTGGGGATCTTCAGCTTC |
| *Gapdh* | AGAAGACTGTGGATGGCCCCTC | GATGACCTTGCCCACAGCCTT |

**Supplementary Data 5. Gene list for the "Matrix Remodeling" custom gene set.** Complete list of 113 genes comprising the "Matrix Remodeling" custom library used for GSEA. The list includes Gene IDs, Symbols, Log2 Fold Changes, and adjusted p-values derived from the microfluidics-engineered organoids vs native lung tissue differential expression analysis. Provided as a `.csv` file.

**Supplementary Data 6. Gene list for the "Immune Response" custom gene set.** Complete list of 145 genes comprising the "Immune Response" custom library used for GSEA. The list includes Gene IDs, Symbols, Log2 Fold Changes, and adjusted p-values derived from the microfluidics-engineered organoids vs native lung tissue differential expression analysis. Provided as a `.csv` file.

**Supplementary Data 7. Gene list for the "Lung Organoid" custom gene set.** Complete list of 56 genes comprising the "Lung Organoid" custom library used for GSEA. The list includes Gene IDs, Symbols, Log2 Fold Changes, and adjusted p-values derived from the microfluidics-engineered organoids vs native lung tissue differential expression analysis. Provided as a `.csv` file.
